# Strong intracellular signal inactivation produces sharper and more robust signaling from cell membrane to nucleus

**DOI:** 10.1101/2020.01.16.909333

**Authors:** Jingwei Ma, Myan Do, Mark. A. Le Gros, Charles S. Peskin, Carolyn A. Larabell, Yoichiro Mori, Samuel A. Isaacson

## Abstract

For a chemical signal to propagate across a cell, it must navigate a tortuous environment involving a variety of organelle barriers. In this work we study mathematical models for a basic chemical signal, the arrival times at the nuclear membrane of proteins that are activated at the cell membrane and diffuse throughout the cytosol. Organelle surfaces within human B cells are reconstructed from soft X-ray tomographic images, and modeled as reflecting barriers to the molecules’ diffusion. We show that signal inactivation sharpens signals, reducing variability in the arrival time at the nuclear membrane. Inactivation can also compensate for an observed slowdown in signal propagation induced by the presence of organelle barriers, leading to arrival times at the nuclear membrane that are comparable to models in which the cytosol is treated as an open, empty region. In the limit of strong signal inactivation this is achieved by filtering out molecules that traverse non-geodesic paths.

## I. INTRODUCTION

Spatial dynamics can play a critical role in the successful functioning of cellular signaling processes, where as basic a property as cell shape can significantly influence the behavior of signaling pathways [1, 2]. Idealized one-dimensional [3], spherical [2, 4, 5] or planar [6] geometries are commonly used in mathematical models of the cell, with the cytosol represented as an empty region of fluid [1–3]. Despite the simplicity of the representation of the plasma membrane and/or cytosolic space, the study of spatial signaling dynamics within mathematical models has provided key insights into the function of many biological pathways, including cyclic AMP signaling in neurons [1], T cell synapse formation through T cell receptor signaling [6], B cell activation through kinase-receptor interactions [4], and general protein kinase signaling [2, 3, 5]. For example, changes in idealized cell shapes can induce significant changes in the timing of signal propagation and the size of concentration gradients across the cytosol [2].

In modeling signal propagation from the cell membrane to the nucleus, a further challenge arises from the crowded, spatially heterogeneous nature of the cytosolic space [7]. In this work we investigate the question of how spatial heterogeneity arising from organelle barriers, as illustrated in Fig. 1b, might influence the propagation of signals from the cell membrane to the nuclear membrane. We consider the simplest possible model for signal propagation from the cell membrane to the nucleus, the release of a one or more activated proteins from the inner cell membrane, and their diffusion throughout the cytosol until they first reach the nuclear membrane. As the classical picture of signal propagation to the nucleus typically involves large pathways of many chemically reacting molecules (such as the MAPK pathway [3]), this model may seem overly simplified. However, a number of proteins are known to be activated at the cell membrane and then directly translocate to the nucleus [8, 9]. For example, in Notch signaling the extracellular domain of Notch receptor can interact with ligands, leading to release of NICD (Notch intracellular domain) from the plasma membrane into the cytosol. NICD then translocates to the nucleus where it can regulate gene transcription [8, 9]. More generally, studying signals that correspond to the diffusive propagation from cell membrane to nucleus of individual proteins provides a first step towards understanding how cellular substructure might influence the dynamics of more complicated signaling pathways.

**FIG. 1:**
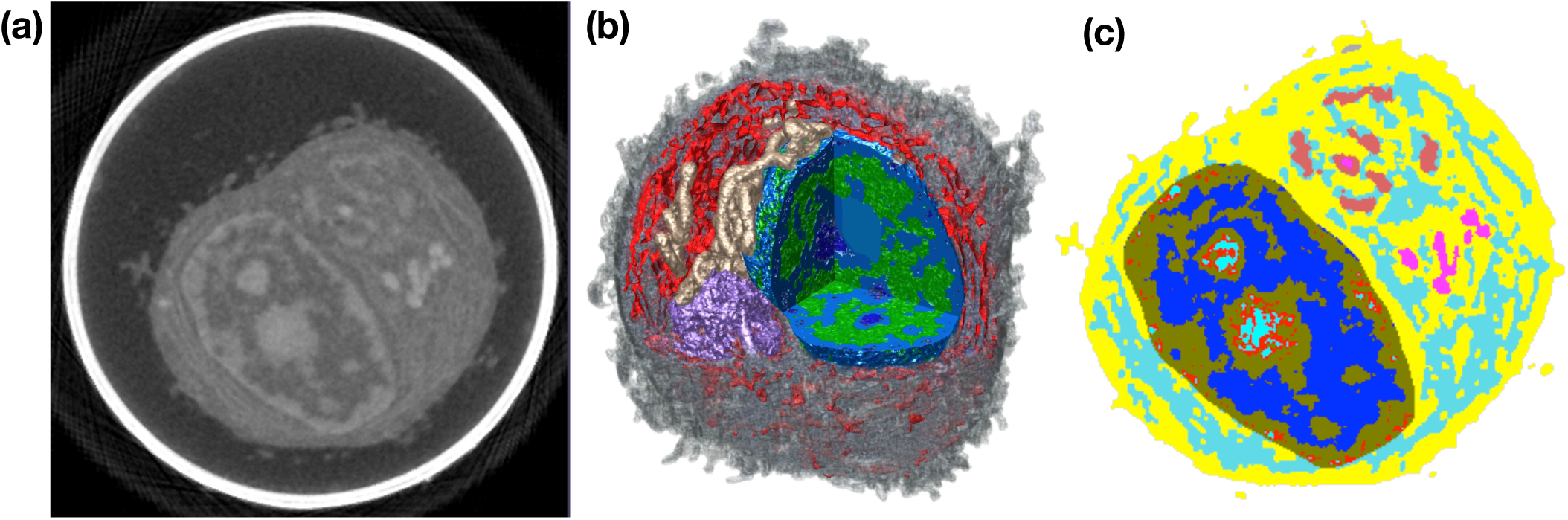
Soft X-ray tomography (SXT) imaging of human B cells. **(a)** One 2D image plane within a 3D SXT reconstruction of a B cell. The corresponding 3D reconstruction is subsequently labeled as Bcell1 in simulations. Pixel intensity corresponds to linear absorption coefficient (LAC), a measure of the local density of organic material [17, 18]. Larger LAC values are shown in lighter colors. The bright white band corresponds to the glass capillary in which the cryo-preserved cell was contained. **(b)** 3D SXT reconstruction of a human B cell with cutaway to show segmented organelles: heterochromatin (blue), euchromatin (green), mitochondria (beige), Golgi (purple) and endoplasmic reticulum (ER) (red). Bulk cytosol is shown in gray, with the cell membrane given by the outer boundary of the cytosol. In our mathematical model, the nucleus, *N*, is given by the set of voxels with labels corresponding to components of the nucleus (e.g. euchromatin and heterochromatin in this image). Cytosol, *C*, is given by voxels rendered in gray, while all other (colored) voxels outside the nucleus are labeled as organelles, *O*. **(c)** Organelle label field values for voxels within the cell in the image plane shown in (a). Here free cytosolic space corresponds to the regions in yellow, and voxels outside the cell are not shown.

Using segmented reconstructions of organelle geometry obtained by soft X-ray tomography (SXT) imaging, we study how the presence of organelle barriers modifies the time needed for diffusing molecules to reach the nucleus in comparison to the time required within an empty cytosol. As signaling molecules diffusing through the cytosol can not persist indefinitely, we next investigate how signal inactivation might influence the search process. This creates a competition where the diffusing signal may be inactivated or degraded prior to reaching the nuclear membrane. We study how the strength of signal inactivation can modulate statistics of the first passage time (FPT) for an individual molecule to reach the nucleus, conditional on it reaching the nucleus before inactivation. It is shown that if the total signal (i.e. number of molecules) that ultimately reach the nucleus is held constant, increasing the inactivation rate leads to signal sharpening. We also find that signal inactivation can provide robustness to the presence of organelle barriers, significantly reducing the difference between the average arrival time of molecules that successfully reach the nucleus in geometries containing organelle barriers, from the time in geometries containing an empty cytosol.

We note that our studies focus on statistics of the time required for the diffusing protein to reach the nucleus. In the case that there is no inactivation, so that the protein simply diffuses until reaching the nucleus, this is an example of a classical diffusion-limited first passage time problem [10]. First passage time problems are widely used in the study of chemical reactions [11, 12], with a variety of asymptotic results and exact solution techniques when the target site is small or a basic geometrical shape such as a sphere [13–16].

## II. MATHEMATICAL MODEL

We consider the time required for a protein to diffuse from the cell membrane to the nuclear membrane. Let *N* denote the nucleus of the cell, with *∂N* denoting the nuclear membrane. Similarly, we let *C* denote the cytosol of the cell, with *∂C* denoting the cell membrane. We assume the cytosol may be filled with a collection of closed subvolumes corresponding to organelles, denoted by *O*, with boundary surfaces *∂O*. Fig 1a shows a slice plane through a 3D soft X-ray tomography (SXT) reconstruction of a human B cell illustrating such geometries, with Fig. 1b showing a 3D reconstruction identifying the nucleus, cytosolic organelles, and the cytosol.

We assume a molecule is initially activated at the cell membrane, and diffuses throughout the cytosolic space until it first reaches the nuclear membrane. Both the cell membrane and organelle surfaces are assumed to be reflecting barriers to the molecule’s diffusion. Denote by *D* = 10(*µ*m)^2^s^−1^ the diffusivity of the molecule, and by *p*(***x***, *t*) the probability density the molecule is located at position ***x*** within *C* at time *t*. ***η***(***x***) will denote the unit outward normal to a surface at ***x***. *p*(***x***, *t*) then satisfies the diffusion equation

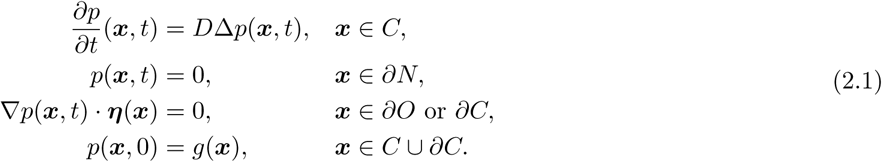

Note, in the following we assume the initial position of the molecule is located on the inner surface of the cell membrane, so that *g*(***x***) is zero away from *∂C*. The Dirichlet boundary condition on *∂N* in (2.1) encodes that the protein is instantly absorbed upon reaching the nuclear membrane, allowing us to study statistics of diffusing protein’s arrival time at the nuclear membrane.

Let *T* denote the random time at which the protein first reaches the nuclear membrane surface. The survival probability that the protein has not yet reached *∂N* at time *t* is then given by

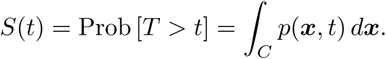

The corresponding probability per time the molecule reaches *∂N* is the probability density function (pdf)

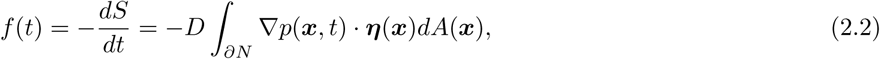

where *dA*(***x***) denotes the surface area measure at ***x*** ∈ *∂N*. Knowing *f*(*t*), we can calculate statistics of *T*, using that the average of a function *w*(*T*), denoted by 𝔼[*w*(*T*)], is defined by

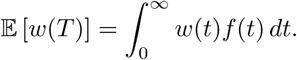

Our representations of cellular geometry are derived from 3D SXT reconstructions, see Materials and Methods, for which the label field identifying organelles is provided as a Cartesian grid of cubes with mesh-width *h*, see Fig. 1. To simulate the time required for the protein to traverse the cytosol we therefore discretize (2.1) onto this grid, generating a system of ODEs we solve numerically. Let *C*_*h*_ denote the collection of mesh voxels that are labeled as being cytosol, with *N*_*h*_ those that are labeled as being within the nucleus, and *O*_*h*_ those within organelles. We label the individual voxels within the cytosol by 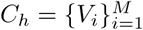, and let *𝒩*(*V*_*i*_; *C*_*h*_) denote the indices of the subset of the six Cartesian grid nearest-neighbors of voxel *V*_*i*_ that are *within* the cytosol. *𝒩*(*V*_*i*_; *N*_*h*_) will similarly denote the indices of the subset of the six Cartesian grid nearest-neighbors of *V*_*i*_ that are within the nucleus. For ***x***_*i*_ denoting the centroid of voxel *V*_*i*_, we let *p*_*h*_(***x***_*i*_, *t*) ≈ *p*(***x***_*i*_, *t*). *p*_*h*_ then satisfies the semi-discrete diffusion equation that

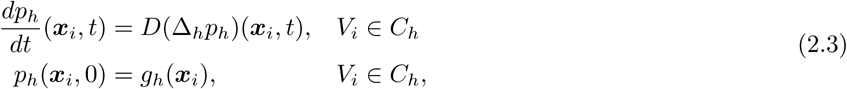

where the discrete Laplacian is defined by

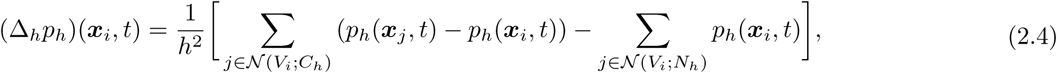

and *g*_*h*_(***x***_*i*_) denotes the initial condition in the semi-discrete model.

This semi-discrete model corresponds to approximating the continuous Brownian motion of the particle in *C* by a continuous-time random walk of the molecule hopping between nearest-neighbor voxels of *C*_*h*_.

If we denote by *T*_*h*_ the corresponding random time for the protein to first reach a voxel that is labeled as being within the nucleus, we have the corresponding survival probability,

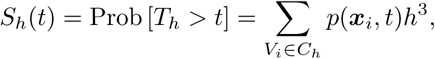

with analogous definitions for the pdf *f*_*h*_(*t*) and averages, 𝔼[*w*(*T*_*h*_)], as above.

In the remainder, unless stated otherwise time will be reported in units of seconds, and distance in units of *µ*m.

## III. ORGANELLE BARRIERS SLOW THE PROPAGATION OF A SIGNAL FROM THE CELL MEMBRANE TO NUCLEUS, WHILE INCREASING VARIABILITY IN ARRIVAL TIME FOR SIGNALS INITIATED AT DIFFERENT LOCATIONS

We begin by numerically solving (2.3) to investigate how the presence of organelles as reflecting barriers influences statistics of the time required for the diffusing protein to reach the nuclear membrane. Let *∂C*_*h*_ denote the collection of voxels within the free cytosol, *C*_*h*_, that border the exterior of the cell, with |*∂C*_*h*_| denoting the volume of this set of voxels. Note, this collection of voxels corresponds to a thin region of cytosol bordering the cell membrane. In the semi-discrete model, we will approximate starting the protein uniformly distributed on the inner surface of the cell membrane by starting the protein uniformly within the volume *∂C*_*h*_. Then

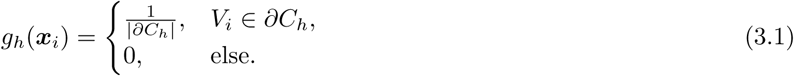

In Fig. 2a we show the survival probability *S*_*h*_(*t*) from Bcell1, the reconstruction shown in Fig. 1 (results from two additional cell reconstructions, labeled Bcell2 and Bcell3, are shown in SI Figures SI1 and SI2). We consider three cases, the physiological data where voxels corresponding to organelles within the cytosol are inaccessible (labeled “physiological”), a modified geometry where voxels corresponding to the endoplasmic reticulum (ER) are added back into the collection of cytosolic voxels the protein can diffuse through (labeled “no ER”), and a modified geometry where all voxels within cytosolic organelles are added back into the collection of cytosolic voxels the protein can diffuse through (labeled “no organelles”). This latter geometry corresponds to the cytosol filling all space between the cell membrane and the nuclear membrane. In Fig. 2a we observe that the presence of organelle barriers dramatically increases the time required for the protein to reach the nuclear membrane (shifting the survival probability curve upwards), with the primary contribution to this shift arising from the barrier provided by the ER. Table I shows that the corresponding mean and median times to reach the cell membrane change similarly. For Bcell1, the presence of the ER as a barrier accounts for most of the the time required to reach the nucleus; removing the ER decreases the median of *T*_*h*_ by almost a factor of three.

**TABLE I:** Statistics of *T*_*h*_, the random time to reach the nucleus in Bcell1. The diffusing molecule is assumed to initially be randomly distributed on the cell membrane, *∂C*_*h*_. Here STD denotes standard deviation and CV denotes the coefficient of variation (the standard deviation divided by the mean). Values in parenthesis denote the ratio of the physiological value to the corresponding no ER or no organelle values. See Table SI1 for statistics in Bcells 2 and 3.

|  | Physiological | No ER | No Organelles |
| --- | --- | --- | --- |
| Bcell1 Mean | 0.7070 | 0.2721 (2.6) | 0.2499 (2.8) |
| Bcell1 Median | 0.4054 | 0.1393 (2.9) | 0.1335 (3.0) |
| Bcell1 STD | 0.8472 | 0.3561 | 0.3173 |
| Bcell1 CV | 1.1983 | 1.3086 | 1.2695 |

**FIG. 2:**
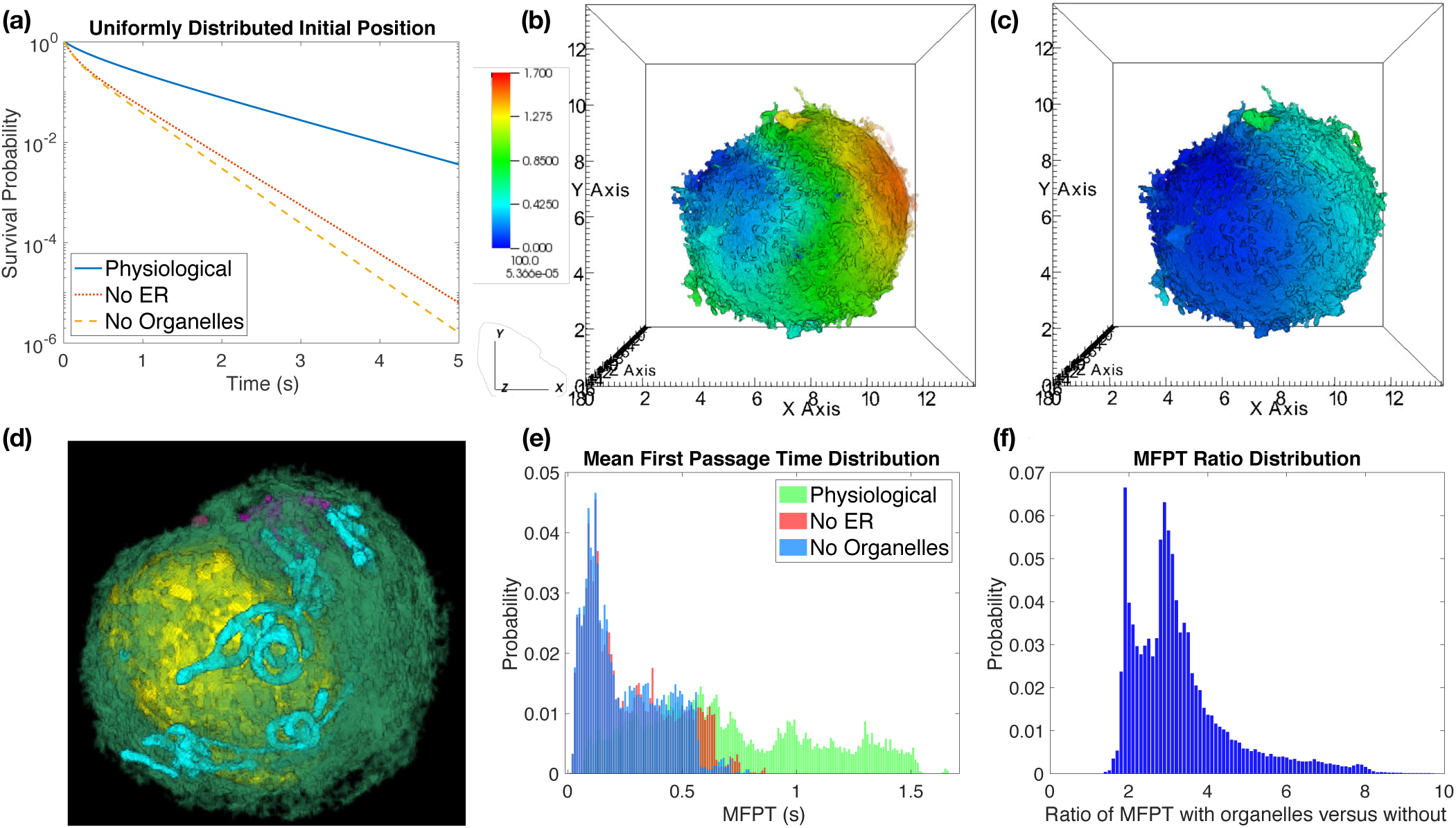
The presence of organelles as diffusive barriers increases the time required for a diffusing (signaling) molecule to traverse from the cell membrane to the nuclear membrane. **(a)** Survival probability, *S*_*h*_(*t*), when the diffusing molecule is started uniformly distributed within a thin region, *∂C*_*h*_, of cytosol bordering the inner surface of the cell membrane (3.1). **(b)** Mean first passage time (MFPT) *u*(***x***_*i*_) from each voxel within *∂C*_*h*_ to reach the nuclear membrane in the “physiological” case that organelles are present as diffusive barriers. Colorbar gives the MFPT values in seconds, spatial units are *µ*m. **(c)** Corresponding MFPTs in the “no organelles” case that the molecules can freely diffuse everywhere between the cell and nuclear membranes. Color scale is the same as (b). **(d)** Volume rendering of the organelles in Bcell1, with the cell in the same orientation as in (b) and (c) (but zoomed in). Note, the ER rendering (green) is attenuated to make other organelles more apparent, and the cell membrane is not shown. Nucleus is in yellow, mitochondria in cyan, and the Golgi in purple. **(e)** Distributions of mean first passage times (MFPTs), 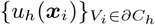, starting from the same thin region of cytosolic voxels bordering the cell membrane as in (b) and (c). Note, here the distribution is over the voxels within the region, illustrating how starting at different initial positions can lead to variation in the MFPT. For the “No ER” case we use the analogous region when just the ER is removed. See (3.2) for definition of the MFPTs *u*_*h*_(***x***_*i*_). Bin width is.01 (seconds). **(f)** Distribution of the ratios of the corresponding “Physiological” to “No Organelles” MFPTS from (e). This illustrates when starting from each individual voxel bordering the cell membrane, how much organelle barriers increase the MFPT to reach the nucleus from that voxel. Bin width is.1. Note, almost all locations have a ratio of two or more, showing that organelle barriers significantly increase the time required to reach the nuclear membrane from most initial positions. SI Figures SI1 and SI2 show similar results for Bcell2 and Bcell3 respectively.

In Figs. 2b-e we examine how the time to reach the nucleus varies when the diffusing molecule is started at different points on the cell membrane. Let *u*(***x***) denote the mean first passage time (MFPT) to diffuse from ***x*** ∈ *C* to the nuclear membrane. *u*(***x***) then satisfies [19]

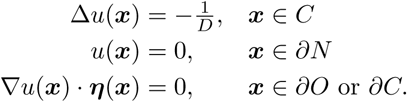

In practice, we solve a discretized version of this PDE that gives the corresponding MFPTs on our Cartesian grid arising from the imaging data. Let *u*_*h*_(***x***_*i*_) denote the MFPT to reach the nucleus from ***x***_*i*_, which satisfies the linear system of equations

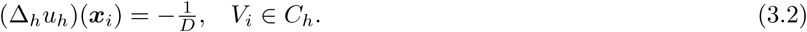

Fig. 2b plots *u*_*h*_(***x***_*i*_) over the cytosolic voxels bordering the cell membrane (*∂C*_*h*_) in the physiological case, while Fig. 2c shows the case with no organelles (i.e. an empty cytosol). We see that the presence of organelles *significantly* slows the MFPT to the nucleus for most points bordering the cell membrane. Not surprisingly, locations closest to the nucleus (left side) generally have smaller MFPTs than locations far from the nucleus (right side). Fig. 2e shows that the distribution of MFPTs, 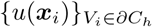, across the cytosolic voxels bordering the cell membrane is much flatter and broader when organelles are present as barriers (green, physiological case) in comparison to an empty cytosol (purple, no organelles case). Moreover, examining the ratio of these MFPTs in the physiological case to the no organelle case, Fig 2f, we find that at almost all locations the presence of organelle barriers increases the MFPT by a factor of two or more.

In conclusion, we observe that organelle barriers can substantially hinder the diffusion of molecules across the cytosol, significantly increasing the time required to reach the nuclear membrane, and increasing the variability of this time *over cytosolic voxels bordering the cell membrane* when comparing signals initiated at different points (Fig. 2f). While our discussion has focused on Bcell1, we observe similar qualitative behavior in Bcell2 and Bcell3, see SI Figures SI1 and SI2.

## IV. INACTIVATION FILTERS OUT MOLECULES UNDERGOING LONGER SEARCHES, REDUCING VARIABILITY IN SIGNAL ARRIVAL TIME

Activated signaling molecules cannot diffuse throughout the cytosol of cells searching for the nuclear membrane indefinitely. Whether by degradation mechanisms, or inactivation mechanisms (such as phosphorylation or dephosphorylation), cellular signals will eventually be terminated. From the perspective of a diffusing signaling molecule this creates a competition between the search for the nuclear membrane and the inactivation process. We now examine how the interplay between these two processes can modulate the timing at which activated signals reach the cell membrane.

We consider the simplest possible mechanism for modeling signal inactivation, assuming the diffusing molecule can now also be inactivated with probability per time *λ*. Let *p*_*λ*_(***x***, *t*) denote the probability density the diffusing molecule is still activated and within the cytosol at time *t. p*_*λ*_ then satisfies

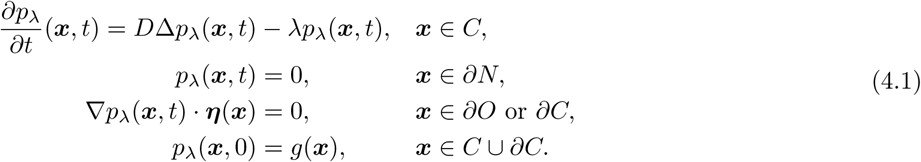

Note that *p*_*λ*_(***x***, *t*) = *e*^−*λt*^*p*(***x***, *t*), so that *p*_0_(***x***, *t*) = *p*(***x***, *t*), the solution to the diffusion equation (2.1).

We are interested in statistics of the exit time through the nuclear membrane, *T*_*λ*_, conditioned on the protein actually reaching the nuclear membrane before inactivation (i.e. the event that *T*_*λ*_ < ∞). The probability per time that the diffusing molecule reaches the nuclear membrane at time *t* is then

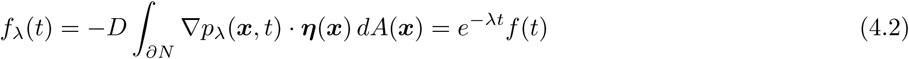

where *f* (*t*) = *f*_0_(*t*) denotes the probability per time to reach the nuclear membrane in the absence of degradation, given by (2.2). With these definitions, the probability the molecule reaches the nuclear membrane before inactivation is

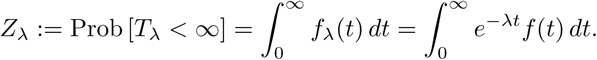

Denoting the conditional cumulative distribution function (CDF) of *T*_*λ*_ by

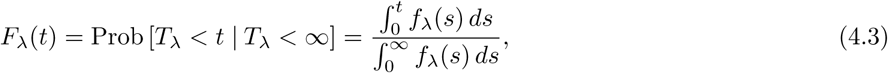

in SI Section SI1 we prove the following results

### Theorem IV.1.

*For all fixed t* > 0 *and λ* ≥ 0, *Z*_*λ*_(*t*) *is a strictly decreasing function of λ, and F*_*λ*_(*t*) *is a strictly increasing function of λ*.

This result gives several immediate corollaries, including that

### Corollary IV.1.

*Both the conditional MFPT*, ⟨*T*_*λ*_⟩ := 𝔼[*T*_*λ*_ | *T*_*λ*_ < ∞], *and the conditional median first passage time*, 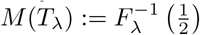, *are strictly decreasing with respect to λ*.

That ⟨*T*_*λ*_⟩ is decreasing in *λ* was also shown in [20] for probability density functions with the factored form *e*^−*λt*^*g*(*t*). Theorem IV.1 and Corollary IV.1 together demonstrate that as the inactivation rate *λ* is increased, the time for a molecule to reach the nucleus, *conditioned* on the molecule actually reaching the nucleus, decreases. The probability any individual molecule actually reaches the nucleus, *Z*_*λ*_, also decreases as *λ* increases. In this way strong signal inactivation will filter out molecules undergoing longer diffusive searches.

To explore how increasing the inactivation rate *λ* influences statistics of the time to reach the nucleus, we now study a semi-discrete model defined on the meshes representing the B cell geometries, and corresponding to a spatial discretization of (4.1). Let *p*_*λ,h*_(***x***_*i*_, *t*) ≈ *p*_*λ*_(***x***_*i*_, *t*) denote the probability density that the diffusing molecule is located at ***x***_*i*_ at time *t*, then

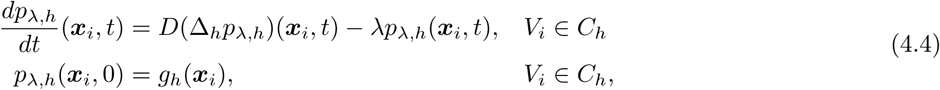

where *p*_*λ,h*_(***x***_*i*_, *t*) = *e*^−*λt*^*p*_*h*_(***x***_*i*_, *t*). Similarly, *f*_*λ,h*_(*t*) = *e*^−*λt*^*f*_*h*_(*t*), so that the probability the diffusing molecule reaches the nucleus is given by

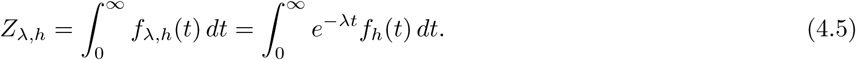

For *T*_*λ,h*_ the random time at which the nucleus is reached, the conditional MFPT to reach the nucleus is then

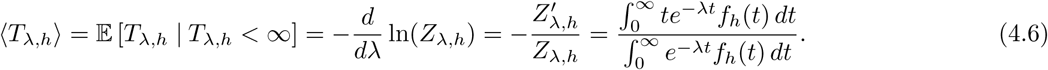

In Figure 3 we consider statistics of *T*_*λ,h*_ when the diffusing molecule is initially placed randomly on the cell membrane (i.e. the uniform initial condition (3.1)). Fig. 3a illustrates Corollary IV.1, showing that for each cell ⟨*T*_*λ,h*_⟩ is strictly decreasing as *λ* is increased. Similarly, Fig. 3c illustrates Theorem IV.1, showing that the probability the molecule reaches the nucleus, *Z*_*λ,h*_, is strictly decreasing as *λ* increases. In Fig. 3b we examine the conditional variance of *T*_*λ,h*_, defined by

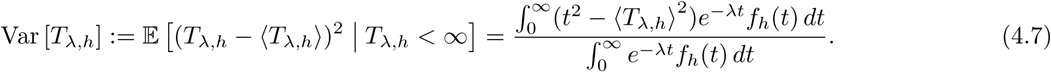

**FIG. 3:**
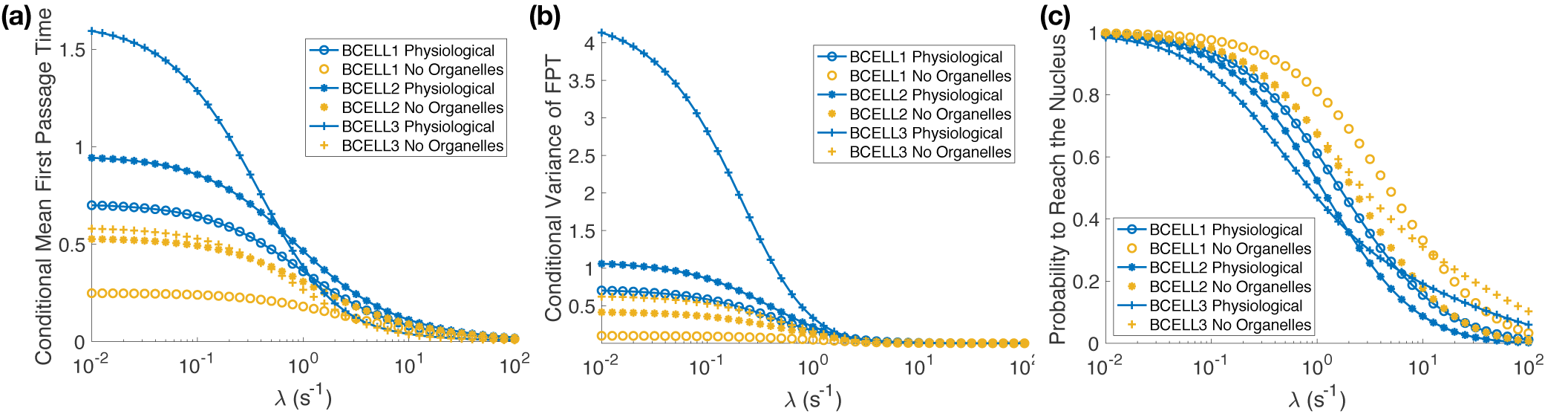
Signal inactivation filters out molecules undergoing longer diffusive searches, reducing both the average time and variance in the time at which a molecule reaches nucleus, conditional on the molecule reaching the nucleus before inactivation. The figures show statistics of the conditional first passage time, *T*_*λ,h*_, to reach the nucleus when the diffusing molecule is started randomly on the cell membrane (i.e. uniformly distributed, see (3.1)), and the molecule can be inactivated with rate *λ*. **(a)** The conditional mean first passage time (MFPT), ⟨*T*_*λ,h*_⟩ (4.6). In all cases we see that ⟨*T*_*λ,h*_⟩ is strictly decreasing as *λ* increases, illustrating Corollary IV.1. Fig. SI4 shows an expanded range of *λ* values, with a logarithmic scale on the *y*-axis. **(b)** The conditional variance of *T*_*λ,h*_, given by (4.7), is decreasing as *λ* increases. **(c)** The probability that the diffusing molecule reaches the nucleus, *Z*_*λ,h*_, is strictly decreasing as *λ* increases, illustrating Theorem IV.1.

In each B cell the conditional variance is strictly decreasing. In SI Figures SI5, SI6 and SI7 we show that similar results hold when the diffusing molecule’s initial position is more localized. Here the molecule is initially placed randomly within small patches of the cell membrane, see SI Section SI2 for details.

## V. INACTIVATION CAN SHARPEN THE SIGNAL REACHING THE NUCLEAR MEMBRANE

To understand how inactivation can affect signal propagation, we investigate how the signal reaching the nucleus changes as the inactivation rate *λ* is increased, but the number of molecules reaching the nucleus is held fixed. By fixing the number of molecules (i.e. total signal) that ultimately reach the nucleus, we can investigate how inactivation influences signal timing without modulating the total signal strength. Note, to fix the total signal reaching the nucleus requires that an increasing number of signaling molecules be released from the cell membrane as *λ* increases.

Consider a deterministic version of (4.4). Assume *N*_0_ molecules are initially uniformly distributed across the interior of the cell, and let *u*_*h*_(***x***_*i*_, *t*) denote the (deterministic) concentration of molecules located at ***x***_*i*_ at time *t*. We assume *u*_*h*_ has units of number per (*µ*m)^3^. *u*_*h*_ then also satisfies (4.4), but with the initial condition

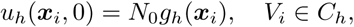

so that *u*_*h*_(***x***_*i*_, *t*) = *N*_0_*p*_*λ,h*_(***x***_*i*_, *t*). The number of molecules per time that successfully reach the nucleus is given by the total flux of *u*_*h*_ into the nucleus, *N*_0_*f*_*λ,h*_(*t*). Similarly, the total number of molecules to successfully reach the nucleus is

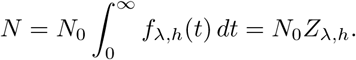

We define the signal reaching the nucleus to be the number of molecules per time that reach the nucleus, given that we assume *N* molecules overall arrive. *N*_0_ is therefore chosen so as to keep *N* fixed as the inactivation rate is varied, so that

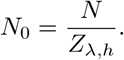

With this choice, the signal, i.e. number of molecules per time, reaching the nuclear membrane is then 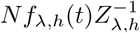. In Fig. 4 we plot the signal reaching the nucleus in Bcell1 as the inactivation rate is increased. SI Fig. SI8 shows the corresponding signals reaching the nucleus in Bcell2 and Bcell3. We see a clear sharpening effect as *λ* increases, with molecules arriving within an earlier and more localized time window. In this context we can interpret increasing activation as speeding up the arrival of the signal at the nuclear membrane.

**FIG. 4:**
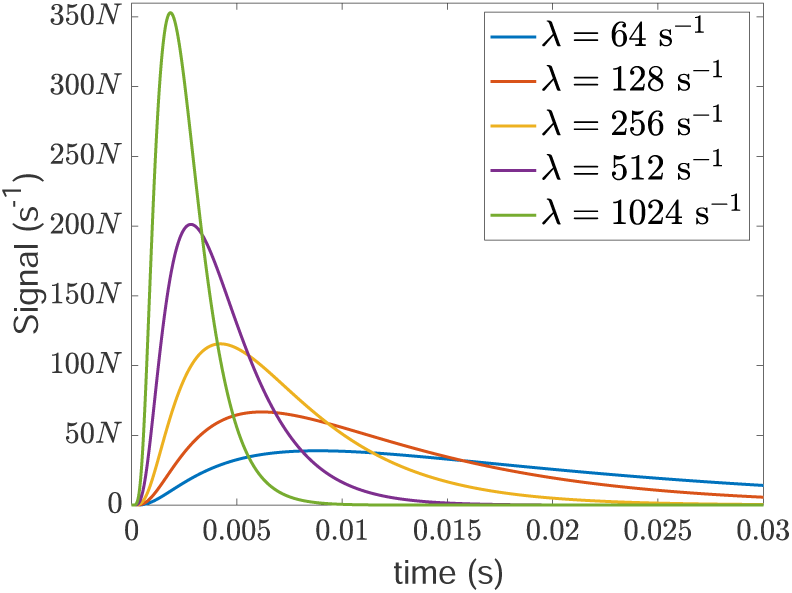
The signal in Bcell1 that successfully reaches the nuclear membrane is sharpened as the inactivation rate, *λ*, is increased. Here signal denotes the expected rate of arrival of signaling molecules at the nuclear membrane when the number of arriving molecules overall is *N*. The expected rate of arrival is plotted as a function of the time that has elapsed since the signaling molecules were released uniformly distributed across the interior of the cell membrane. Note that the total number of *arriving* molecules is being held constant in the results plotted here, and this requires that more signaling molecules be released when *λ* is greater. This is achieved by choosing the total number of molecules that are released initially as 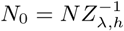. As explained in Section V, in a deterministic model with this initial condition, the signal corresponds to the flux (number of molecules per time) successfully reaching the nucleus (given by 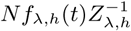). For the single-particle stochastic model (4.4), *N*0 = *N* = 1 and the signal corresponds to the first passage time density to reach the nucleus, conditional on the molecule arriving before inactivation (given by 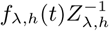). A similar signal sharpening effect is observed in Bcell2 and Bcell3, see SI Fig. SI8.

While the deterministic model shows the window in which the molecules arrive becomes smaller as inactivation increases, the single-particle stochastic model (4.4) allows us to see how much variation one would have in the number of molecules that successfully reach the nucleus. We again assume that *N*_0_ signaling molecules are activated uniformly on the interior of the cell membrane, and that the molecules’ dynamics are *completely independent*. The number of molecules that reach the nucleus would then be a binomial random variable, **N** ∼ *B*(*N*_0_, *Z*_*λ,h*_), in *N*_0_ with parameter *Z*_*λ,h*_. The average number of molecules to reach the nucleus would be 𝔼 [**N**] = *N*_0_*Z*_*λ,h*_, while the coefficient of variation in the number of molecules to reach the nucleus is

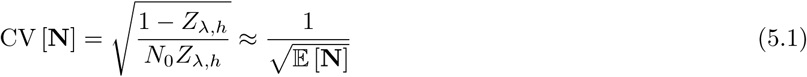

for *λ* large. Here we have used that the probability to reach the nucleus, *Z*_*λ,h*_ approaches zero as *λ* → ∞, see the next section, and approximated the square root in the numerator by the leading-order term of its Taylor series expansion about *Z*_*λ,h*_ = 0. Keeping *N*_0_*Z*_*λ,h*_ fixed as the inactivation rate is increased then preserves the expected number of molecules to reach the nucleus. Moreover, (5.1) demonstrates that the relative variation in the number of molecules that reach the nucleus will be small if the average number of molecules that reach the nucleus, 𝔼 [**N**], is sufficiently large. By modulating both the inactivation rate and the number of signaling molecules released at the cell membrane, a cell can then tune both how localized the signal is in time, and the noisiness in the number of molecules that successfully reach the nuclear membrane.

## VI. INACTIVATION CAN PROVIDE ROBUSTNESS WITH RESPECT TO CELLULAR SUBSTRUCTURE IN THE TIME FOR A SIGNAL TO REACH THE NUCLEUS

In Fig. 5a we plot the ratio of ⟨*T*_*λ,h*_⟩ in the physiological case to the no organelles case. For very small values of the inactivation rate the figure demonstrates that the presence of organelles can significantly increase the time required for one diffusing molecule to reach the nucleus. In contrast, as *λ* increases, for each B cell we see that the ratio decreases to a value close to one. That is, strong signal inactivation seems to be able to buffer out the effect of cellular geometry. This comes at the cost of a significantly decreased probability any individual signaling molecule will reach the nucleus.

**FIG. 5:**
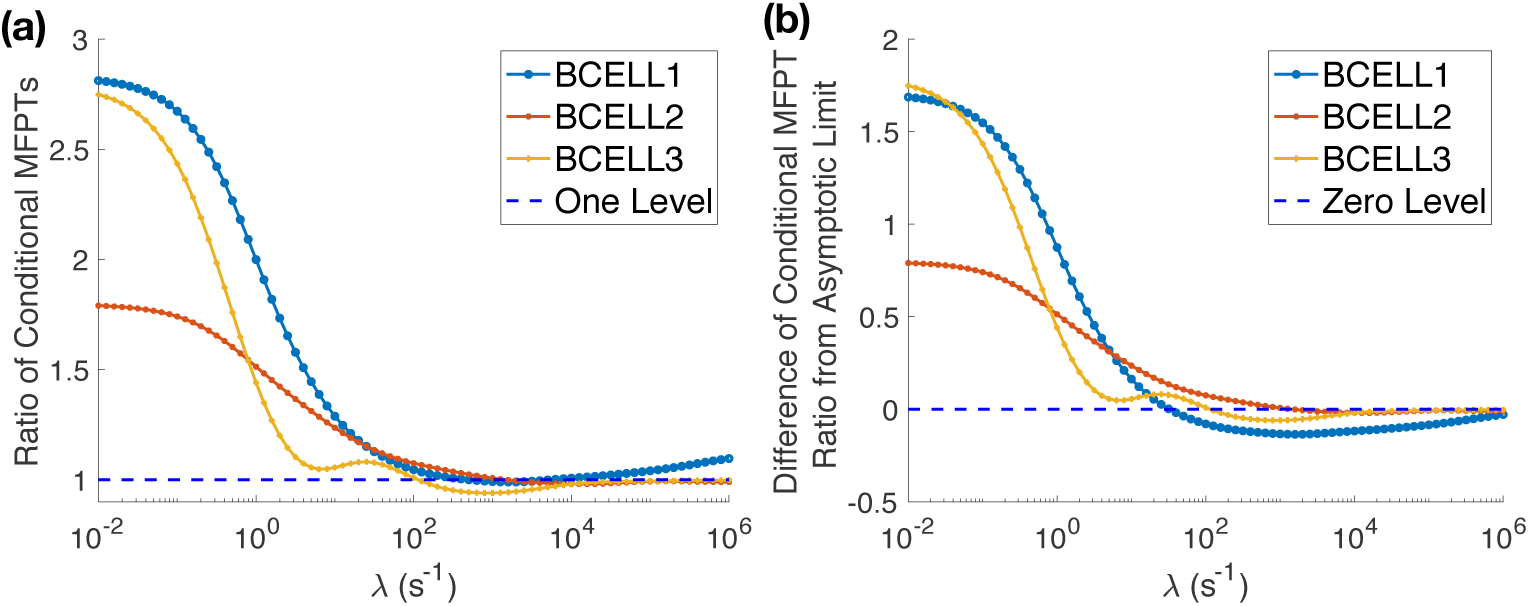
Strong signal inactivation can buffer out the effects of cellular substructure on the time to find the nucleus. **(a)** The ratio of the conditional mean first passage time (MFPT) to reach the nucleus, *T*_*λ,h*_, in the physiological case to the conditional MFPT in the no organelles case decreases significantly from its initial value as *λ* increases. For each cell the ratio approaches a number close to one, indicating that strong signal inactivation can completely buffer out the effect of cellular substructure on the time to find the nucleus. **(b)** Difference of the ratio of ⟨*T*_*λ,h*_⟩ shown in (a) from its asymptotic limit (6.4). Note, (b) demonstrates that the slight increase above one for the ratio (6.4) in Bcell1 is just the approach to its asymptotic limit, 1.125. The ratios (6.4) for Bcell2 and Bcell3 both converge to 1 in (a).

These simulations illustrate that the ratio of the MFPTs between the physiological and no organelle cases is decreased for sufficiently strong signal inactivation. To understand the limit to how much strong signal inactivation can buffer out the effect of organelle barriers in our model, we now examine the large *λ* asymptotic expansion of the conditional MFPT, ⟨*T*_*λ,h*_⟩. Our goal is to derive an explicit formula for the asymptotic limit of ⟨*T*_*λ,h*_⟩ as *λ* → ∞ that illustrates the role of the geometry of the cytosolic space. Our derivation demonstrates how the effect of geometry on the limiting conditional MFPT arises. Readers interested solely in the derived formula may skip ahead to (6.3).

By (4.6), knowing the asymptotic behavior of *Z*_*λ,h*_ as *λ* → ∞ would allow us to calculate the behavior of ⟨*T*_*λ,h*_⟩. In turn, the behavior of *Z*_*λ,h*_ can be calculated from the integral representation (4.5). This will be determined by the short-time behavior of *f*_*h*_(*t*) due to the rapid decay of the exponential for large *λ*. We therefore begin by examining the behavior of *f*_*h*_ as *t* → 0. We can estimate this short-time behavior by direct Taylor series expansion using a matrix exponential representation for the evolution operator, i.e.

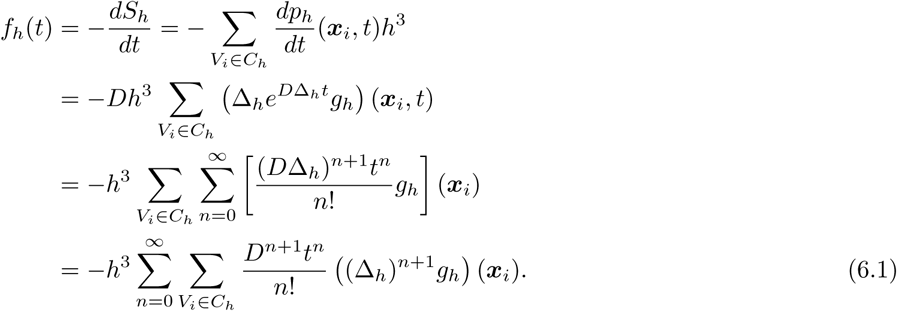

To simplify this expression we make use of the relationship between powers of the discrete Laplacian and geodesic (nearest-neighbor) graph distances.

Recall our assumption that *g*_*h*_(***x***_*i*_) = 0 for all ***x***_*i*_ ∉ *∂C*_*h*_, and denote by 𝒢_*h*_ ⊂ *∂C*_*h*_ the set of voxels in which *g*_*h*_(***x***_*i*_) ≠ 0 (i.e. the support of *g*_*h*_). If the particle is started randomly placed within the voxels bordering the cell membrane then 𝒢_*h*_ = *∂C*_*h*_, whereas if the particle is initially started at a fixed point, ***x***_*i*_, then 𝒢_*h*_ = {***x***_*i*_}. Given a set of voxels 𝒱 ⊂ *C*_*h*_, we define *d*(𝒱, *N*_*h*_) to be the shortest (integer) graph distance along a nearest-neighbor path from each voxel in 𝒱 to first reach a voxel in *N*_*h*_. Here by nearest-neighbor we mean the six nearest-neighbors to a given voxel, two from each of the ***x, y*** and ***z*** directions. For example, if no voxel in 𝒱 is within *N*_*h*_, but *some* voxel in 𝒱 has a nearest neighbor that is within *N*_*h*_, then *d*(𝒱, *N*_*h*_) = 1.

It is from the powers of the discrete Laplacian in (6.1) that the role of cytosolic geometry in the short-time behavior of *f*_*h*_(*t*) arises, ultimately dictating the large *λ* behavior of ⟨*T*_*λ,h*_⟩. As shown in SI Lemma.1, the {*V*_*i*_ ∈ *C*_*h*_ | (Δ_*h*_)^*k*^*g*_*h*_(***x***_*i*_) ≠ 0 will contain no voxels *bordering* the nucleus until *k* = *d*(𝒢_*h*_, *N*_*h*_) − 1. For any smaller *k*, one additional application of the discrete Laplacian then simply moves probability mass within the cytosol. As such, mass is conserved and we have the following result which is proven in SI Section SI1

### Theorem VI.1.

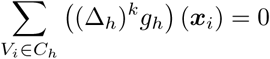

*for* 1 ≤ *k* ≤ *d*(𝒢_*h*_, *N*_*h*_) − 1.

With *d*_*g*_ = *d*(𝒢_*h*_, *N*_*h*_), the theorem implies that (6.1) can be simplified to

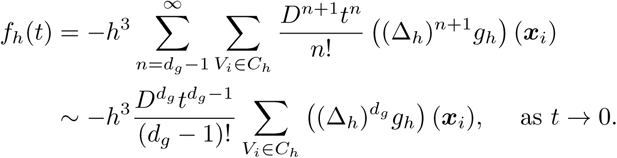

Assuming that *d*_*g*_ > 1, we obtain the corresponding estimate for *Z*_*λ,h*_ as *λ* → ∞ by

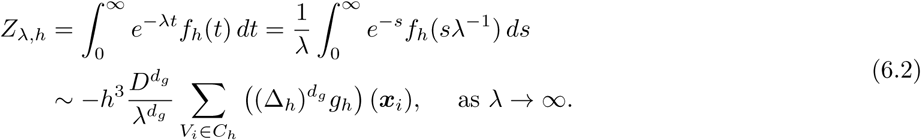

In SI Theorem.1 we prove this asymptotic formula holds. Taking logarithmic derivatives, we find that

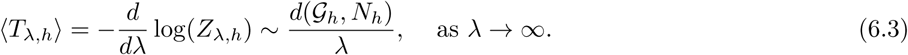

In SI Figure SI4 we show the convergence of ⟨*T*_*λ,h*_⟩ to this asymptotic formula as *λ* → ∞.

Let *d*(𝒢_*h*_, *N*_*h*_)_phys_ denote the distance from 𝒢_*h*_ to the nucleus in the physiological case, with *d*(𝒢_*h*_, *N*_*h*_)_n.o._ the distance in the no organelle case. Define ⟨*T*_*λ,h*_⟩_phys_ and ⟨*T*_*λ,h*_⟩_n.o._ analogously. The ratio of the conditional MFPTs then approaches

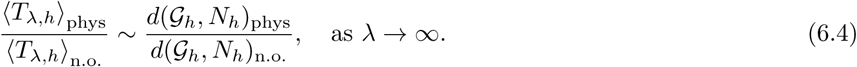

That is, how much the effect of geometry on the search time can be buffered out by strong inactivation in our model is essentially controlled by how the shortest path (nearest-neighbor) graph distance from the initial set the particle can be placed in to the nucleus changes between the physiological and no organelle cases. In particular, since the voxels within the cytosol in the physiological case are always a strict subset of those in the no organelles case, we see the ratio is always at least one (in the limit).

In Figure 5b we plot the difference between the ratio of the conditional MFPTs and the derived asymptotic limit in (6.4). We see that for each cell the asymptotic limit is approached as *λ* → ∞, but that the approach is not always monotonic. In particular, the asymptotic limit (6.4) does not appear to be a rigorous lower bound for how much the effect of geometry can be buffered out over all possible inactivation rates.

If the diffusing molecule is started at a fixed location, ***x***_*i*_, we obtain

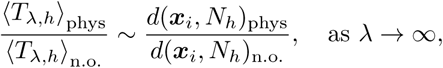

the ratio of the shortest graph (nearest-neighbor) distances from ***x***_*i*_ to the nucleus in the two cases. In particular, if the shortest path distance from ***x***_*i*_ to the nucleus is the same in both cases, we find that the effect of organelle barriers on the conditional MFPT is completely filtered out in the limit of strong signal inactivation.

In SI Section SI2, we show analogous results to Figure 5 when the diffusing molecule is started randomly within small patches of the cell membrane. We see similar qualitative behavior for statistics of *T*_*λ,h*_, and for the ratio of ⟨*T*_*λ,h*_⟩ in the physiological to no organelles cases. Note, however, that we observe a variation in how much the effect of geometry can be buffered out as the patch of cell membrane where the signal is initiated moves about.

## VII. DISCUSSION

Our results demonstrate that organelle barriers to the molecular diffusion of signaling molecules can significantly slow the propagation of a signal from the cell membrane to the nucleus. Such barriers also increase the variability in the distribution of times to reach the nucleus for signals activated at different localized portions of the cell membrane. Strong signal inactivation provides one potential mechanism to both buffer out the effect of organelle barriers, and to reduce variability in the time at which signals reach the nucleus. Mechanisms to reduce such variability may be needed to ensure robust functioning of pathways that involve pulsatile responses. For example, the relative expression of the pituitary hormones LH and FSH is controlled by the pulse frequency of extracellular GnRH ligands [21]. Sufficient variability in processing such signals might lead to improper expression levels through misidentification of the pulse frequency.

As shown in Section V, under the constraint that the expected number of molecules to reach the nucleus should be *fixed* at *N*, the inactivation rate can be adjusted provided that the initial number of molecules activated at the inner surface of the cell membrane are varied in a compensating manner. Under these assumptions, Fig. 4 demonstrates that the time for a signal to reach the nuclear membrane can be made arbitrarily small by increasing the inactivation rate. This comes with a clear cost though; increasing the rate of signal inactivation requires increasing numbers of signaling molecules to be activated at the cell membrane to maintain a fixed number of molecules that successfully reach the nucleus.

Our conclusions can be generalized in several ways. First, while we focused on the propagation of a signal between the cell and nuclear membranes, our results should hold more generally for a variety of signal sources and targets within cells. In more general signaling pathways they should also apply to the most downstream signaling component, presuming it is not activated right near the nuclear membrane. Finally, we note that while signaling pathways can involve complicated reaction kinetics throughout the cytosol, it may be that in some cases their overall effect can be approximated as a single signal that propagates throughout the cytosol and is inactivated on some timescale.

### Regime of Model Applicability

It is important to note that the large *λ* asymptotic scaling in (6.3), and convergence to the ratio (6.4), may require relatively large values of *λ* (on the order of *λ* between 10^4^ s^−1^ and 10^6^ s^−1^ for *D* = 10 (*µ*m)^2^s^−1^, see Figure 5b and SI Figure SI4). Molecules that successfully reach the nucleus would on average arrive on time scales of 10^−4^s^−1^ or less, see Figure SI4, which would not necessarily be expected to be physically plausible in a typical mammalian cell. More generally, as *λ* → ∞ these results rely on the (increasingly) short-time behavior of the continuous-time random walk model (4.4). However, both the continuous diffusion model (4.1) and the continuous time random walk model (4.4) become physically unrealistic as models for the very short-time motion of a molecule within a cell. Moreover, the *very* short-time behavior of the semi-discrete model (4.4) and the continuous diffusion model (4.1) would not be expected to agree, since the former only approximates the latter on sufficiently large timescales.

The relative behavior of the two models is illustrated in Fig. SI9 and SI Section SI3. There we compare the analytical PDE solution, when the nuclear membrane and cell membrane are represented as concentric spheres, to the numerical solution of the corresponding semi-discrete model on a Cartesian grid approximation of the cytosolic region between the spheres. We find that for a mesh spacing of *h* = 0.0351*µ*m, comparable to that of our B cell reconstructions, ⟨*T*_*λ*_⟩ and ⟨*T*_*λ,h*_⟩ agree exceptionally well until the asymptotic *λ*^−1^ scaling takes over in the semi-discrete model. Then we see a discrepancy due to the different short-time behavior of the semi-discrete model, with the *λ*^−1^ scaling, and the exact solution to the continuous diffusion PDE, which exhibits a *λ*^−1*/*2^ scaling, see (SI5).

For these reasons the usefulness of understanding the large *λ* asymptotic behavior is not in the predicted scaling of ⟨*T*_*λ,h*_⟩ (6.3), but in the decreasing asymptotic behavior of the conditional MFPT ratio (6.4). This asymptotic limit provides insight into why, on physiological timescales, we observe a decrease in the effect of organelle barriers on signal propagation. Namely, signal inactivation filters out the molecules that would have had to traverse longer paths to get to the nucleus. This reduces differences between the lengths of paths which molecules that reach the nucleus must take in the organelle filled, and organelle empty, cell.

### Conjectures and Open Problems

For the continuous diffusion model (4.1), let 𝒢 denote the set on which *g*(*x*) ≠ 0 (i.e. the support of *g*(*x*)). For example, if the particle is started uniformly on the inner surface of the cell membrane than 𝒢 = *∂C*. We conjecture that the corresponding ratio of conditional MFPTs satisfies

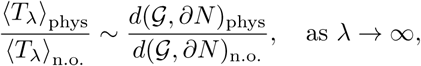

where *d*(𝒢, *∂N*) refers to the shortest path geodesic distance through the cytosol from the signal initiation location, 𝒢, to the nuclear membrane *∂N*. We have obtained partial results to this effect when there are straight line paths from 𝒢 to *∂N* and the principal curvatures of the nuclear membrane satisfy certain constraints, but the general case remains an open problem.

The geodesic distance has recently been suggested to also arise in the context of the first searcher problem. Here one is interested in the average time at which the first of *N* searchers reaches a target as the number of searchers, *N*, becomes large (i.e. *N* → ∞). In [22] it was suggested that, similar to our observations for strong signal inactivation, this limit also filters out all but the shortest paths, with the average time for the first searcher to reach a target scaling like the square of the geodesic distance. An interesting future question would be to understand the interplay of these two problems; i.e. the time required for the first of many searchers to successfully reach a binding target in the presence of strong signal inactivation.

Finally, we note that it is an open question to understand whether spatial signaling pathways [3, 23, 24] involve more general mechanisms for filtering out the effect of spatial heterogeneity within the cytosolic environment. It would be particularly interesting to investigate such questions while also studying the role of two effects that we have not explicitly resolved; crowding between molecules within the cytosol and active transport of signaling molecules to the nuclear membrane. In addition, in this work we considered only the simplest of signaling components: linear inactivation. For many signaling pathways, including BCR signaling in B cells and general protein kinase signaling, inactivation is more appropriately modeled as occurring through a nonlinear interaction with a phosphatase [4, 5]. Such pathways also commonly involve cascades of interactions [3], which could conceivably have additional mechanisms that buffer out the influence of cellular substructure on signal timing. We hope to explore such models in future work.

## VIII. MATERIALS AND METHODS

### Reconstruction of Cellular Substructure

To reconstruct the locations of organelles and membrane surfaces, we made use of soft X-ray tomographic (SXT) imaging of cells. For an overview of SXT imaging, we refer the reader to [18]. In this work we used reconstructions of three human B cells (GM12878 lymphoblastoids) from [25]. The experimental protocol for obtaining these reconstructions was also described in [25]. SXT is similar in concept to medical X-ray CT imaging, but uses soft X-rays in the “water window,” which are absorbed by carbon and nitrogen dense organic matter an order of magnitude more strongly than by water [18]. As the absorption process satisfies the Beer–Lambert law, the measured linear absorption coefficient (LAC) of one voxel of a 3D reconstruction is linearly related to the density of organic material within that voxel [18]. In practice, SXT reconstructions are able to achieve resolutions of 50 nm or less. For all reconstructions used in this work, the underlying voxels were cubes with sides of length 0.03515625 *µ*m. Another advantage of SXT is in the minimal preprocessing of cells that is required before imaging. Cells are cryogenically preserved, but no segmentation, dehydration, or chemical fixation is necessary. Figure 1a shows the reconstructed LAC values from one image plane within a 3D SXT reconstruction of Bcell1.

As discussed in [26], many organelles have different underlying densities of organic material, and therefore attenuate soft X-rays differently. This is reflected in their having different LAC values. Exploiting this property, 3D SXT reconstructions were labeled and segmented in Amira [27], using a combination of Amira’s automated segmentation tools based on LAC values, followed by hand segmentation to refine segmentation boundaries [26]. Each underlying voxel within the 3D SXT reconstruction was labeled as belonging to one of a variety of organelles (heterochromatin, euchromatin, endoplasmic reticulum, mitochondria, Golgi apparatus, bulk cytosol, etc.). Figure 1c shows one plane of the resulting label field.

### Numerical Solution of Semi-discrete Diffusion Equation (2.3)

The semi-discrete diffusion equation (2.3) was solved in PETSc 3.7.7 [28, 29] using the adaptive Runge-Kutta Chebyshev (RKC) method of [30] with both the absolute and relative errors set to 10^−8^. To evaluate the solution, *p*_*h*_(***x***, *t*), at larger times, it was approximated by a truncated eigenvector expansion using all terms with eigenvalues having a magnitude less than one. The corresponding eigenvalues and eigenvectors of the discrete Laplacian (2.4) were calculated in SLEPc 3.7.4 [31] using the Krylov-Schur solver with default parameter values and tolerances. For all simulations the decision to switch from the RKC solver to the eigenvector expansion was made by looking over the interval 1 < *t* < 10 for where the two solutions first differed by an absolute error of less than 10^−5^ and a relative error of less than.01.

To numerically evaluate the integrals defining statistics such as *Z*_*λ,h*_ and ⟨*T*_*λ,h*_⟩, we split them into two pieces. The integral from zero to the time at which the PDE solver switched from the RKC method to the truncated eigenvector expansion, and the integral from this time to infinity. The first integral was evaluated using the cumulative trapezoidal rule at the discretization times used in the RKC method. The second integral was evaluated by analytically integrating the truncated eigenvector expansion. Within these integrals the probability density function for the molecule to reach the nucleus was calculated directly from the flux into voxels of the nucleus,

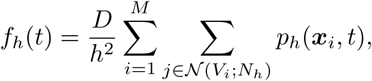

using the numerically computed solutions.

## ACKNOWLEDGMENTS

SAI and JM were supported by National Science Foundation awards DMS-1255408 and DMS-1902854. SAI thanks the Isaac Newton Institute of Mathematical Sciences for hosting him for the Programme on Stochastic Dynamical Systems in Biology while working on this project. During this time SAI was partially supported by a Simons Fellowship of the Isaac Newton Institute. A portion of the reported simulations made use of the Boston University Shared Computing Cluster. Research reported in this publication was conducted at The National Center for X-ray Tomography, located at the Advanced Light Source of Lawrence Berkeley National Laboratory, and was supported by grants from NIH (P41GM103445) and DOE’s Office of Biological and Environmental Research (DE-AC02-5CH11231). MAL and CAL were supported by NIH (U1DA040582) and Chan Zuckerberg Initiative Human Cell Atlas Program.

## SUPPORTING INFORMATION

### SI1. Proofs of Main Text Theorems

In this section we give proofs of the results in Sections IV and VI. We begin with Theorem IV.1 and Corollary IV.1 from Section IV. In our proofs, we do not show here, but make use of, the following basic results on properties of the solutions to (2.1) and (4.1)

1. The probability per time the diffusing molecule reaches the nucleus in the model without degradation, *f* (*t*), is positive for *t* > 0.
2. The conditional survival probability *S*_*λ*_(*t*) = 1 − *F*_*λ*_(*t*) approaches zero as *t* → ∞ sufficiently fast that *tS*_*λ*_(*t*) → 0 and *S*_*λ*_(*t*) is integrable.

#### Theorem IV.1.

*Proof*. Let *λ*_2_ > *λ*_1_ ≥ 0, it is immediate that 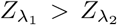. Showing 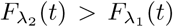 for all *t* > 0 is equivalent to showing

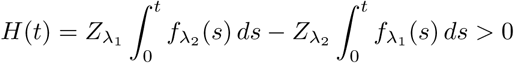

for *t* > 0. Note *H*(0) = 0 and lim_*t*→∞_ *H*(*t*) = 0. We then have that

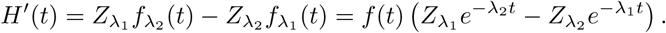

As such, using that *f* (*t*) > 0 for *t* > 0, we have that *H*′(*t*) = 0 if and only if

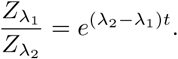

As the left side is greater than one, there is exactly one point, *t*^∗^, where equality can hold for 0 < *t* < ∞. For 0 < *t* < *t*^∗^, *H*′(*t*) > 0, while for *t* > *t*^∗^, *H*′(*t*) < 0. As such, we conclude that *H*(*t*) increases from *H*(0) = 0 to a global max, and then decreases to zero as *t* → ∞, implying that *H*(*t*) > 0 for 0 < *t* < ∞. □

#### Corollary IV.1.

*Both the conditional MFPT*, ⟨*T*_*λ*_⟩ := 𝔼[*T*_*λ*_ | *T*_*λ*_ < ∞], *and the conditional median first passage time*, 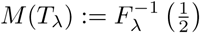, *are strictly decreasing with respect to λ*.

*Proof*. By definition

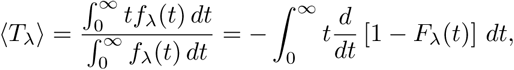

where the conditional CDF, *F*_*λ*_(*t*), is given by (4.3). Note that *F*_*λ*_(*t*) ∈ [0, 1], *F*_*λ*_(0) = 0 and lim_*t*→∞_ *F*_*λ*_(*t*) = 1. Using the assumed integrability of *tS*_*λ*_(*t*) = *t*(1 − *F*_*λ*_(*t*)), and integrating by parts, then gives

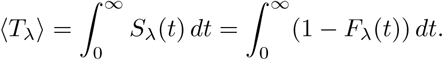

As *F*_*λ*_(*t*) is strictly increasing with respect to *λ* for *t* > 0 and *F*_*λ*_(*t*) > 0 for *t* > 0, this implies that ⟨*T*_*λ*_⟩ is strictly decreasing in *λ*.

To prove the monotonicity of the median, let *λ*_2_ > *λ*_1_ ≥ 0. Then

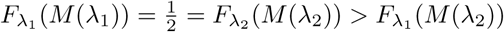

by the monotonicity of *F*_*λ*_(*t*) with respect to *λ*. Since the CDF *F*_*λ*_(*t*) is non-decreasing and continuous in *t*, we conclude that *M* (*λ*_1_) > *M* (*λ*_2_). □

We now prove Theorem VI.1 from Section VI. Let supp *w*_*h*_ denote the set of voxels in *C*_*h*_ where a lattice function *w*_*h*_(***x***_*i*_) is non-zero. We first prove the following lemma, which shows that sup{(Δ_*h*_)^*k*^*g*_*h*_} will contain no voxels *bordering* the nucleus until *k* = *d*(𝒢_*h*_, *N*_*h*_) − 1. For any smaller *k*, one additional application of the discrete Laplacian then simply moves probability mass within the cytosol. As such, mass is conserved and we have the following result

#### Lemma.1.

*Let k* ∈ {0, 1, …, *d*(supp{*g*_*h*_}, *N*_*h*_) − 1}. *Then d* (supp{(Δ_*h*_)^*k*^*g*_*h*_}, *N*_*h*_) = *d*(*g*_*h*_, *N*_*h*_) − *k*.

*Proof*. The case *d*(sup{*g*_*h*_}, *N*_*h*_) = 1 is immediate. Assume *d*(supp{*g*_*h*_}, *N*_*h*_) ≥ 2. Let *k* ≤ *d*(supp *g*_*h*_, *N*_*h*_) − 2, and assume Σ_*k*_ = sup{(Δ_*h*_)^*k*^*g*_*h*_} is the set of nearest neighbor shortest paths of length *k* from supp{*g*_*h*_}. By definition of the discrete Laplacian, Σ_*k*+1_ then contains Σ_*k*_, and all nearest-neighbors of Σ_*k*_ within *C*_*h*_. This corresponds to all nearest-neighbor shortest paths from the voxels within supp{*g*_*h*_} of integer length ≤ *k* + 1, so that *d*(Σ_*k*_, *N*_*h*_) = *d*(Σ_*k*+1_, *N*_*h*_) + 1. Since *d*(Σ_0_, *N*_*h*_) = *d*(supp{*g*_*h*_}, *N*_*h*_) by definition, the theorem then holds by induction. □

The lemma implies that supp{(Δ_*h*_)^*k*^*g*_*h*_} will contain no voxels *bordering* the nucleus until *k* = *d*(supp{*g*_*h*_}, *N*_*h*_) − 1. For any smaller *k*, one additional application of the discrete Laplacian simply moves probability mass within the cytosol, but outside of voxels that border the nucleus. This then implies Theorem VI.1:

#### Theorem VI.1.

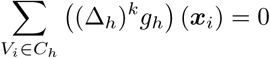

*for* 1 ≤ *k* ≤ *d*(𝒢_*h*_, *N*_*h*_) − 1.

*Proof*. Assume *d*(supp{*g*_*h*_}, *N*_*h*_) ≥ 2. We may then write

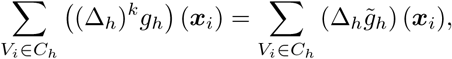

where 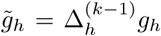. By Lemma.1 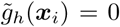 for all ***x***_*i*_ that are nearest-neighbors to voxels within the nucleus. Using (2.4), when acting on 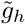 the discrete Laplacian then simplifies to

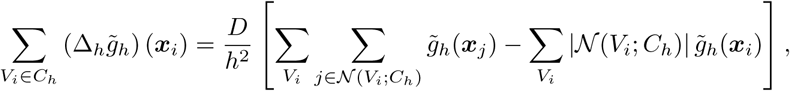

where |*𝒩* (*V*_*i*_; *C*_*h*_)| denotes the number of neighbors of voxel *V*_*i*_ within *C*_*h*_. Reordering the first sum we have that

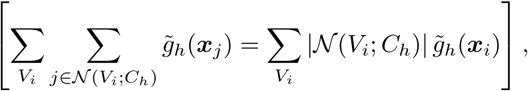

so that

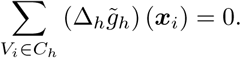

□

In the following theorem we prove the asymptotic behavior of *Z*_*λ,h*_ given in (6.2).

#### Theorem.1.

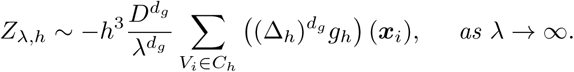

*Proof*. By definition

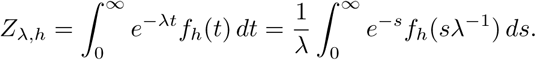

Plugging in the expansion formula for *f*_*h*_(*t*)

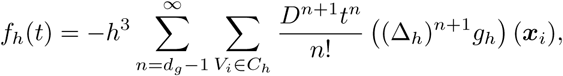

we have that

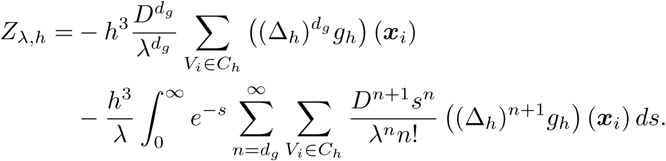

In the last equation, denote the second, remainder term by *I*. We claim 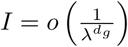 as *λ* → ∞. We have

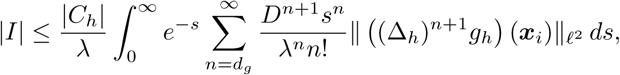

where |*C*_*h*_| denotes the volume of the cytosol and 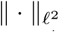 denotes the discrete *ℓ*^2^ norm over *C*_*h*_. Let *σ*_*max*_ label the largest singular value of the discrete Laplacian matrix, Δ_*h*_, then

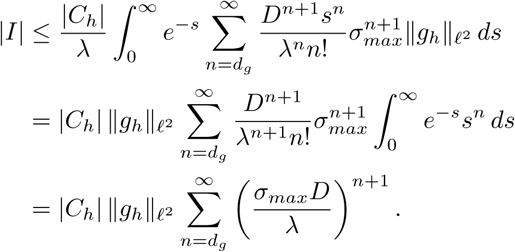

Assuming we take *λ* > *σ*_*max*_*D* large enough the last series is convergent and we have

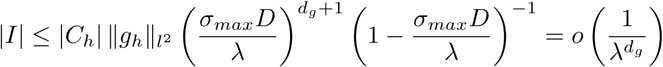

as *λ* → ∞. □

### SI2. Statistics of the time to reach the nucleus with localized initial conditions

To understand how localization of the initiation of signals might influence the time for a signal to reach the nucleus, we also conducted simulations using localized patch initial conditions. This corresponded to the initial condition

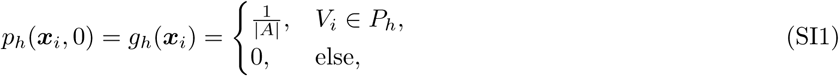

where *P*_*h*_ denotes the set of voxels within a given patch of the cell membrane and *A* the area of the patch.

100 patches were determined for each cell by selecting 100 seed points on the cell membrane of the “physiological” geometries (i.e. cells with all internal organelles present). A 100-bin, equally-spaced histogram for the distribution of MFPTs across the cell membrane was generated from the values of *u*_*h*_(***x***_*i*_), see (3.2). From each bin one seed location was then randomly sampled from the collection of voxels with MFPTs within that bin. About each seed point a patch was constructed by adding all nearest-neighbor voxels of the seed point that were also within the membrane. The procedure was then repeated, adding all nearest-neighbors to previously calculated neighbors. This procedure was then repeated recursively for the newly added voxels until at least 100 voxels were obtained. The final patch then formed a connected graph within the cell membrane containing all *k* nearest neighbors of the seed voxel for some value *k*. In Figure SI3 we show the distribution of patch diameters for the 100 patches sampled for each B cell. Typical final patch sizes were between.3 and.5 *µ*m in diameter.

In Figures SI5, SI6, and SI7 we show statistics of the conditional MFPT to reach the nucleus, *T*_*λ,h*_, for Bcell1, Bcell2 and Bcell3 respectively. In each case we see similar qualitative behavior in the statistics to what we observed for the uniform initial condition used in the main text, see Figures 3 and 5.

### SI3. Comparison of Semi-Discrete Model to Continuous PDE Model: Spherically Symmetrical Case

We now investigate the accuracy in approximating, and differences between, our semi-discrete model and the continuous diffusion equation model. As our cellular reconstructions are given as labeled Cartesian meshes, we do not have an underlying spatially-continuous domain representation with corresponding analytic solution to which we can compare. We therefore instead consider a simpler problem, the spherically symmetrical case. We model the cell as a 3D ball with radius *R* centered at the origin, and model the nucleus as a 3D concentric ball of smaller radius *r*. While this problem is idealized, we will solve it using comparable mesh sizes to the B cell reconstructions, allowing us to characterize how well we resolve the continuous Brownian Motion of molecules when using this resolution in spherical geometries. In the case of continuous Brownian Motion between a spherical nucleus and cell we provide a standard, but self-contained, derivation for *Z*_*λ*_ and ⟨*T*_*λ*_⟩. We note that in the pure-diffusive case such results are well-known [13].

We first consider an initial condition starting from a point ***y*** on the cell membrane. Taking the Laplace transform of (4.1) and denoting 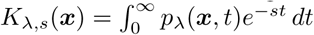, we obtain

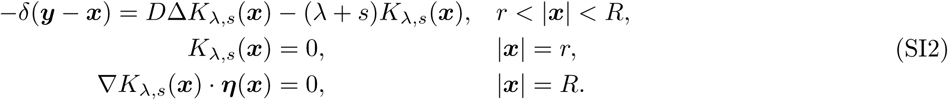

Let *F*_*λ,s*_(***y***) = −*D*∫_*∂N*_ ∇*K*_*λ,s*_(***x***, *t*) · ***η***(***x***) *dA*(***x***). *F*_*λ,s*_(***y***) can be solved from the following PDE:

#### Lemma.2.

*F*_*λ,s*_(***x***) *is a solution to the following boundary value problem:*

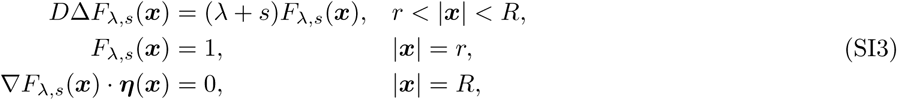

*where* ***η***(***x***) *denotes the unit outward normal to the sphere ∂B*(0, *R*) = {|***x***| = *R*}.

*Proof*. Multiplying (SI3) by *K*_*λ,s*_(***x***) and applying Green’s identity, we obtain

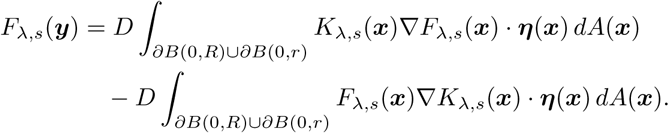

Plugging in the boundary conditions of (SI2) and (SI3), we find *F*_*λ,s*_(***y***) = −*D*∫_*∂N*_ ∇*K*_*λ,s*_(***x***, *t*) · ***η***(***x***) *dA*(***x***). □

Solving (SI3), we obtain

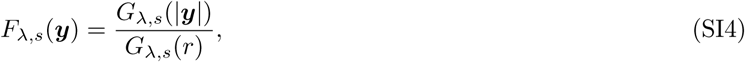

where

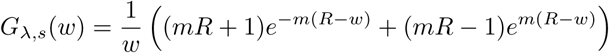

and

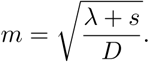

The probability that starting from a point ***y***, the molecule reaches the nuclear membrane before inactivation can be rewritten as

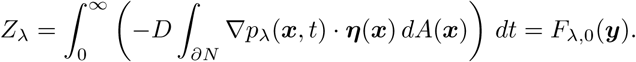

From (SI4), this probability only depends on the length of the initial position, |***y***|. Therefore, the same exit time statistics hold for a uniformly distributed initial condition starting from the cell membrane, i.e. |***y***| = *R*, so that

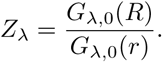

The conditional mean first passage time can be calculated from

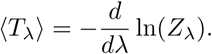

In particular, when *λ* → 0 we obtain

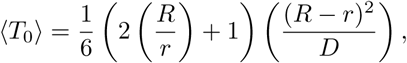

and when *λ* → ∞,

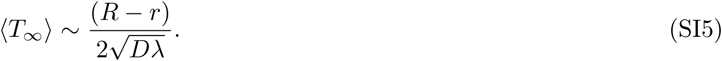

As we mention in the main text Discussion, this result is reflective of the short-time behavior of the signaling molecule’s continuous Brownian motion. Since random walks do not approximate Brownian motions on sufficiently short time scales, as expected the *λ*^−1*/*2^ scaling we obtain is different than the *λ*^−1^ scaling we proved for the semi-discrete model.

For numerical comparison, we generated a cell-centered 3D Cartesian mesh to approximate the spherically symmetrical geometry, where if the center of a voxel is within the ball of radius *r* = 3*µ*m we identified it as being in the nucleus. Likewise, if the center of a voxel is within the ball radius of *R* = 5*µ*m but outside the nucleus, we identified it being in the cytosol. Voxels cut by the sphere of radius *R* = 5*µ*m were identified as belonging to the cell membrane. The mesh width in our simulations was *h* = 0.0351*µ*m, which is comparable to the mesh size for each of the B cells we studied. The numerical solution method described in the Methods section was used for solving the corresponding semi-discrete model with one alteration. We used a slightly coarser absolute error tolerance of 1e-4 and relative error tolerance of.01 for determining the time at which to switch from the Runge-Kutta-Chebyshev method to the truncated eigenvector expansion.

In Figure SI9 we compare the analytical ⟨*T*_*λ*_⟩ given by the logarithmic derivative of *F*_*λ*,0_(*R*) to the numerical solution of the semi-discrete model. We see that the two solutions agree exceptionally well until the large *λ* asymptotic behavior takes over. Both solutions still continue to decrease as *λ* is further increased, but with the different asymptotic scalings discussed above. We also include a second semi-discrete solution, for a smaller mesh width of *h* = 0.0175*µ*m, which used only the RKC method for solving in time. (We found it challenging to calculate the needed eigenvectors on the finer mesh, and examination of the semi-discrete solution in the coarser mesh case showed that the contribution of the eigenvector expansion at longer times to Figure SI9 was minimal. This makes intuitive sense since the short-time behavior dominates in calculating ⟨*T*_*λ,h*_⟩.) We see that the finer mesh results in a larger range of *λ* over which ⟨*T*_*λ,h*_⟩ approximates ⟨*T*_*λ*_⟩ well before the asymptotic behavior takes over.

### SI4. Supplemental Figures and Tables

**FIG. SI1:**
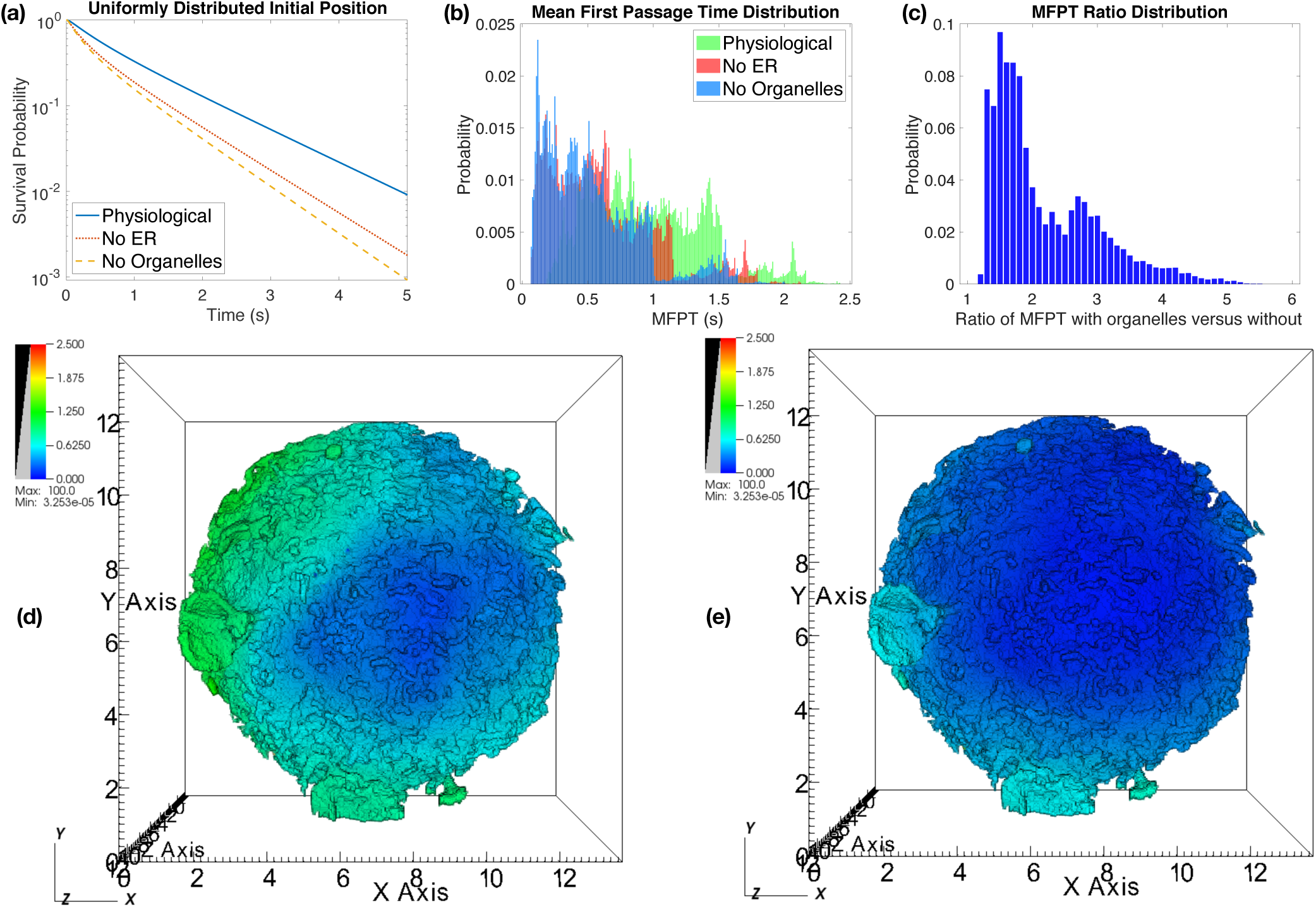
Statistics of MFPT in the absence of signal degradation for Bcell2. See Figure 2 for subfigure information.

**TABLE SI1:** Statistics of *T*_*h*_, the random time to reach the nucleus in the absence of signal degradation in Bcell1, Bcell2 and Bcell3. The diffusing molecule is assumed to initially be randomly distributed on the cell membrane, *∂C*_*h*_. Here STD denotes standard deviation and CV denotes the coefficient of variation (the standard deviation divided by the mean).

|  | Physiological | No ER | No Organelles |
| --- | --- | --- | --- |
| Bcell1 Mean | 0.7070 | 0.2721 | 0.2499 |
| Bcell2 Mean | 0.9540 | 0.6041 | 0.5311 |
| Bcell3 Mean | 1.6360 | 0.6841 | 0.5861 |
| Bcell1 Median | 0.4054 | 0.1393 | 0.1335 |
| Bcell2 Median | 0.5964 | 0.3358 | 0.2941 |
| Bcell3 Median | 0.8842 | 0.3087 | 0.2706 |
| Bcell1 Variance | 0.7178 | 0.1268 | 0.1007 |
| Bcell2 Variance | 1.0850 | 0.5390 | 0.4210 |
| Bcell3 Variance | 4.3352 | 0.9015 | 0.6337 |
| Bcell1 CV | 1.1983 | 1.3086 | 1.2695 |
| Bcell2 CV | 1.0919 | 1.2154 | 1.2218 |
| Bcell3 CV | 1.2726 | 1.3880 | 1.3582 |

**FIG. SI2:**
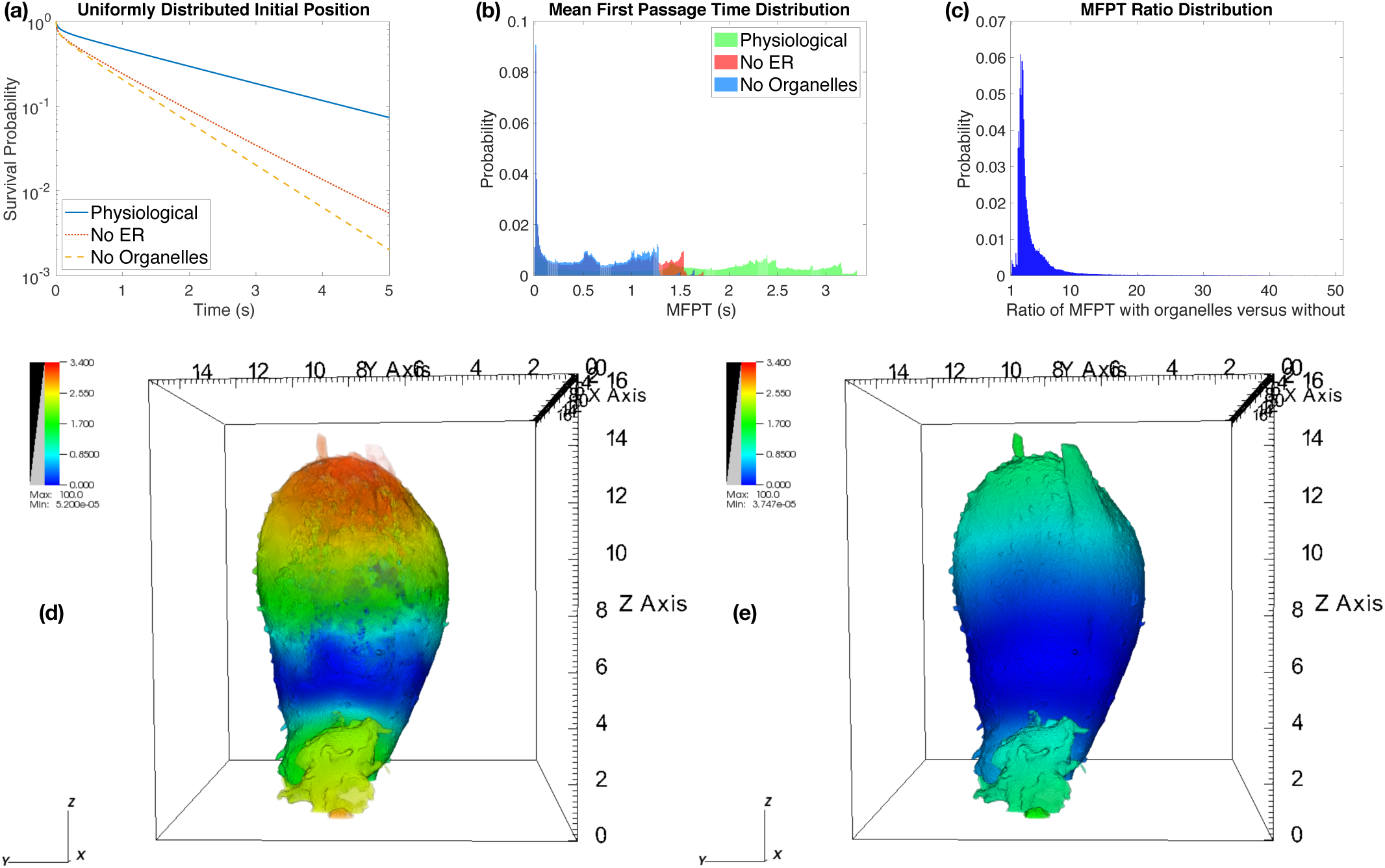
Statistics of MFPT in the absence of signal degradation for Bcell3. See Figure 2 for subfigure information.

**FIG. SI3:**
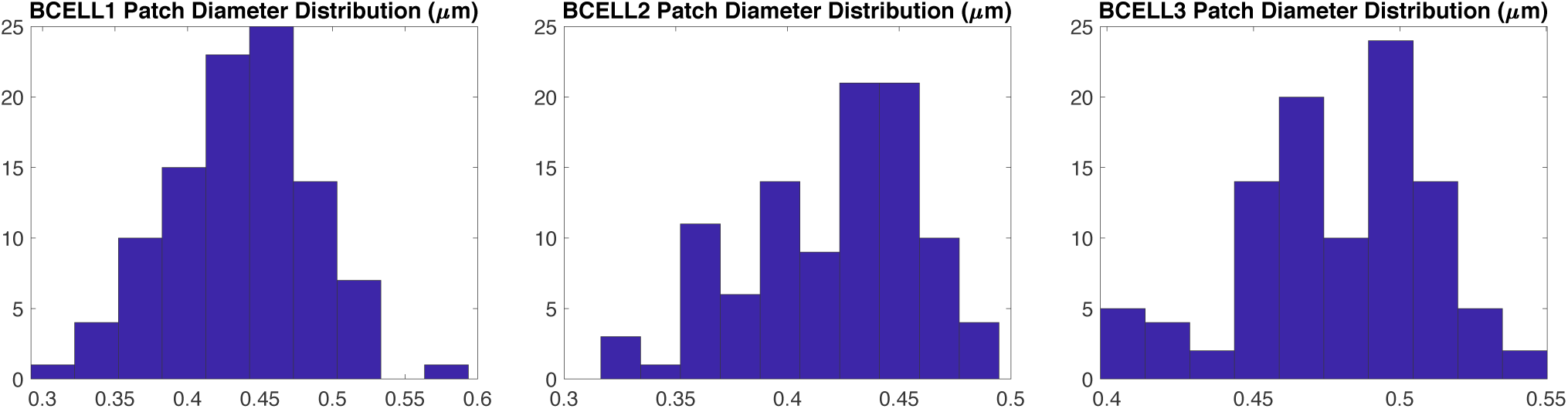
Distribution of patch diameters for the 100 patches in Bcell1, Bcell2 and Bcell3. Here diameter corresponds to the largest Euclidean distance between the center of two voxels within the patch.

**FIG. SI4:**
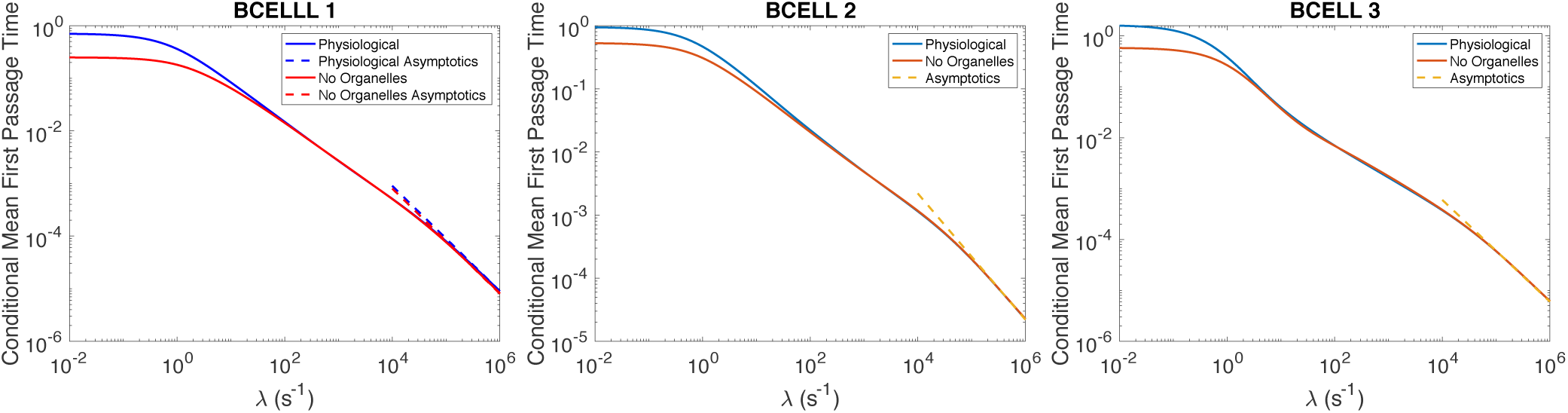
Convergence of ⟨*T*_*λ,h*_⟩ to the asymptotic limit (6.3) as *λ* → ∞ when the molecule is started uniformly on the surface of the cell. In Bcell1, the geodesic distance from the cell membrane to the nucleus is different in the “physiological” and “no organelles” cases, while in Bcell2 and Bcell3 the distance is the same (and so only one asymptotic line is shown).

**FIG. SI5:**
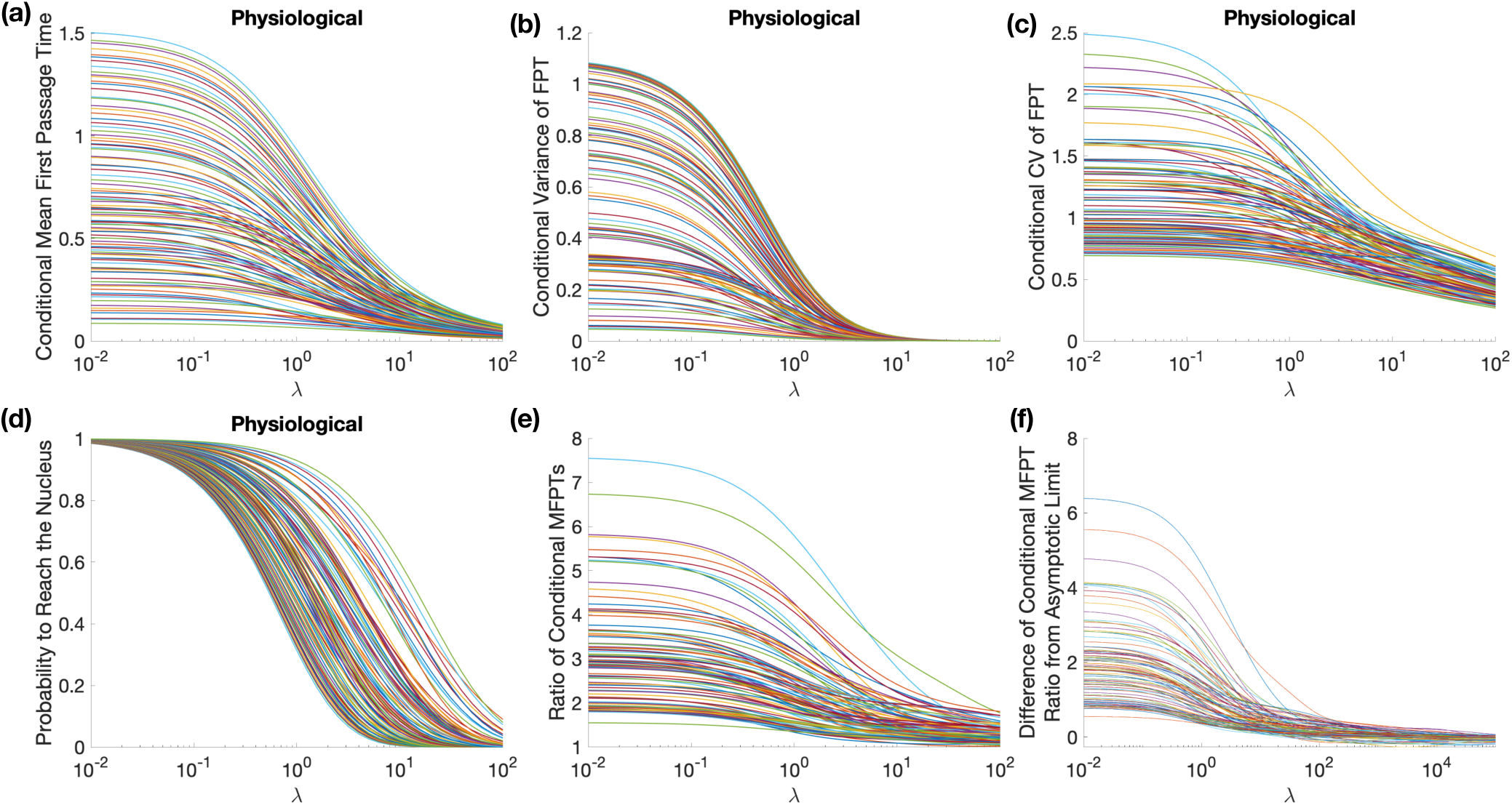
Statistics of the conditional MFPT, *T*_*λ,h*_, for Bcell1 for 100 different patch initial conditions (see Section SI2). (a) through (d) show statistics for the physiological case. (e) shows the ratio of the physiological to no organelle conditional MFPTs, while (f) shows the difference between this ratio and the asymptotic limit.

**FIG. SI6:**
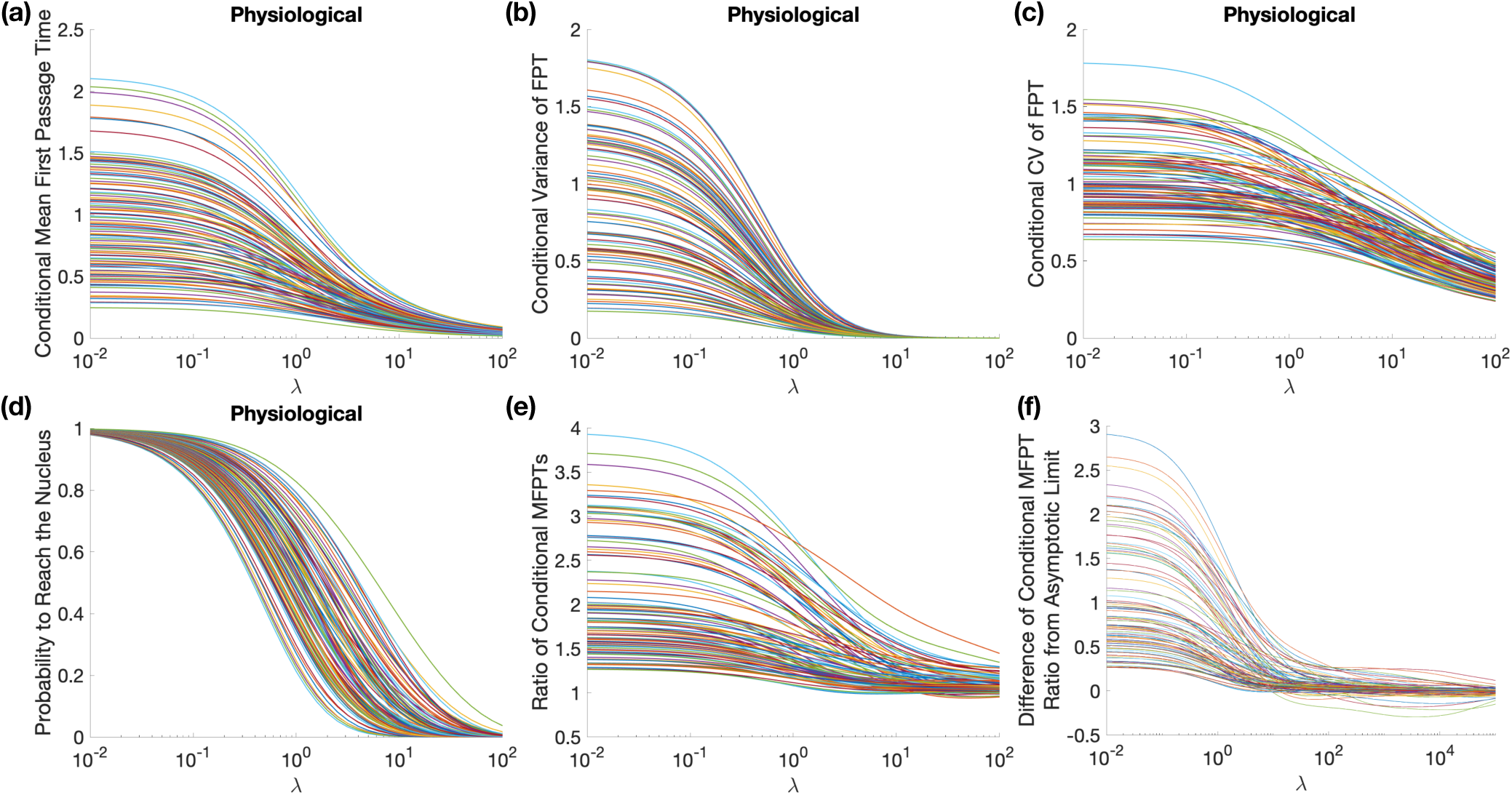
Statistics of the conditional MFPT, *T*_*λ,h*_, for Bcell2 for 100 different patch initial conditions (see Section SI2). (a) through (d) show statistics for the physiological case. (e) shows the ratio of the physiological to no organelle conditional MFPTs, while (f) shows the difference between this ratio and the asymptotic limit.

**FIG. SI7:**
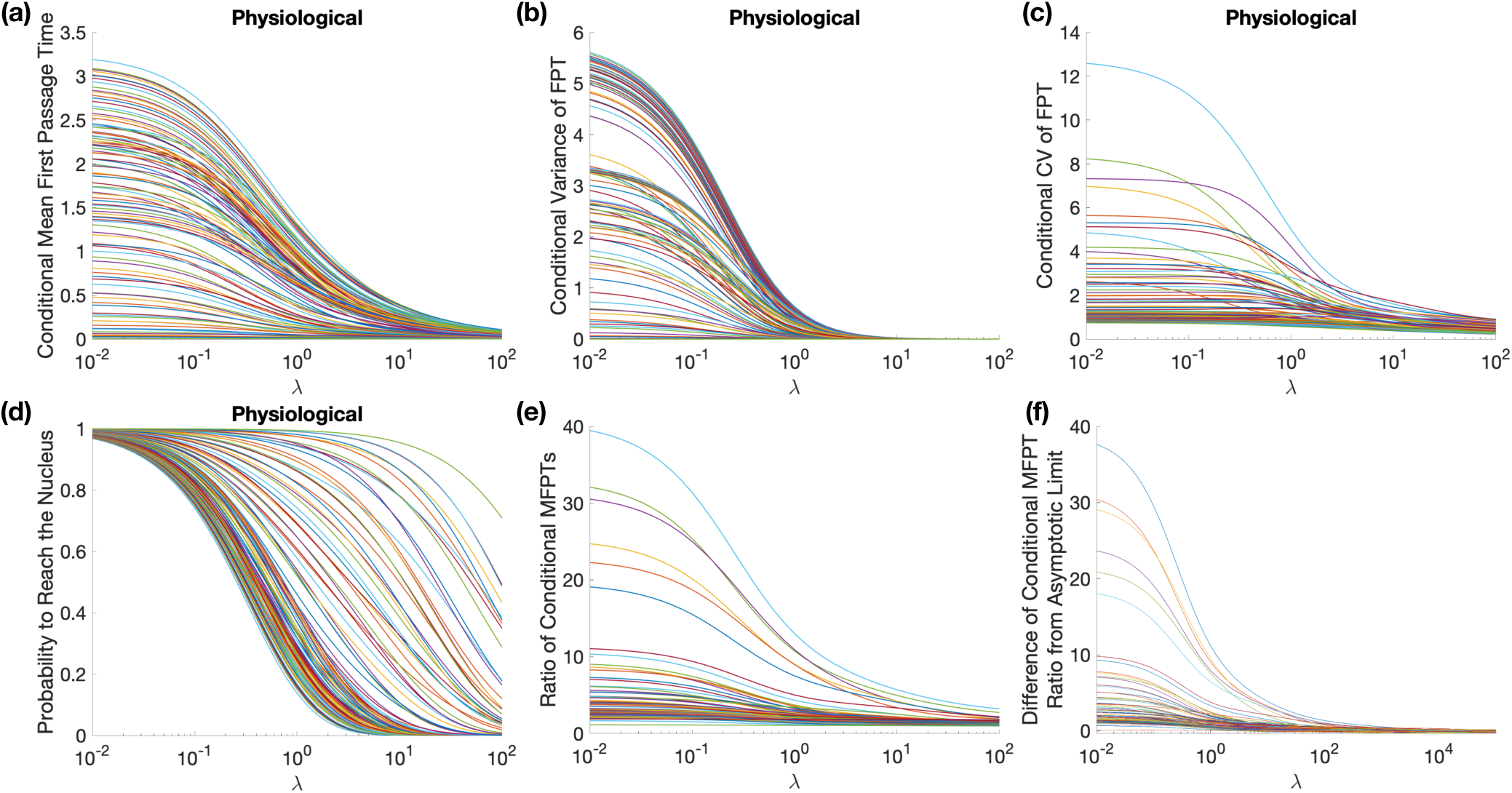
Statistics of the conditional MFPT, *T*_*λ,h*_, for Bcell3 for 100 different patch initial conditions (see Section SI2). (a) through (d) show statistics for the physiological case. (e) shows the ratio of the physiological to no organelle conditional MFPTs, while (f) shows the difference between this ratio and the asymptotic limit.

**FIG. SI8:**
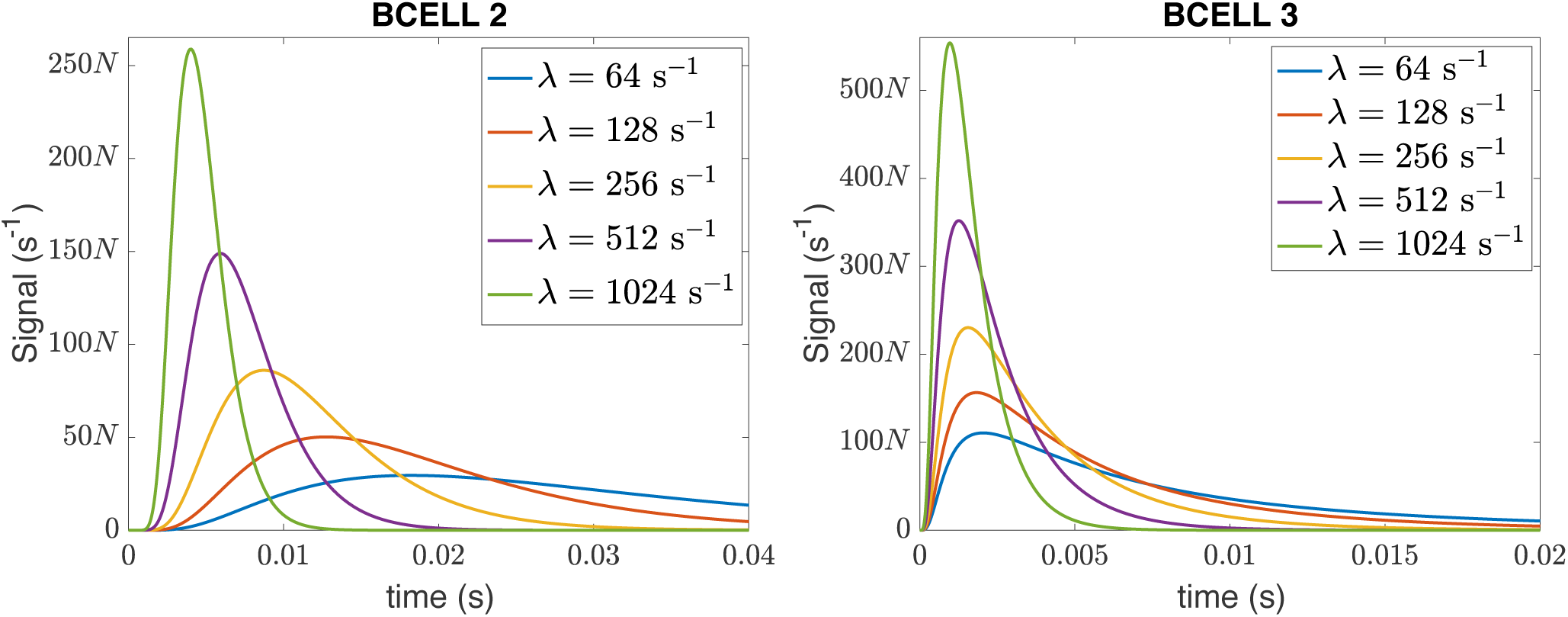
Signal successfully reaching the nucleus in Bcell2 and Bcell3. See Fig. 4 in the main text for details.

**FIG. SI9:**
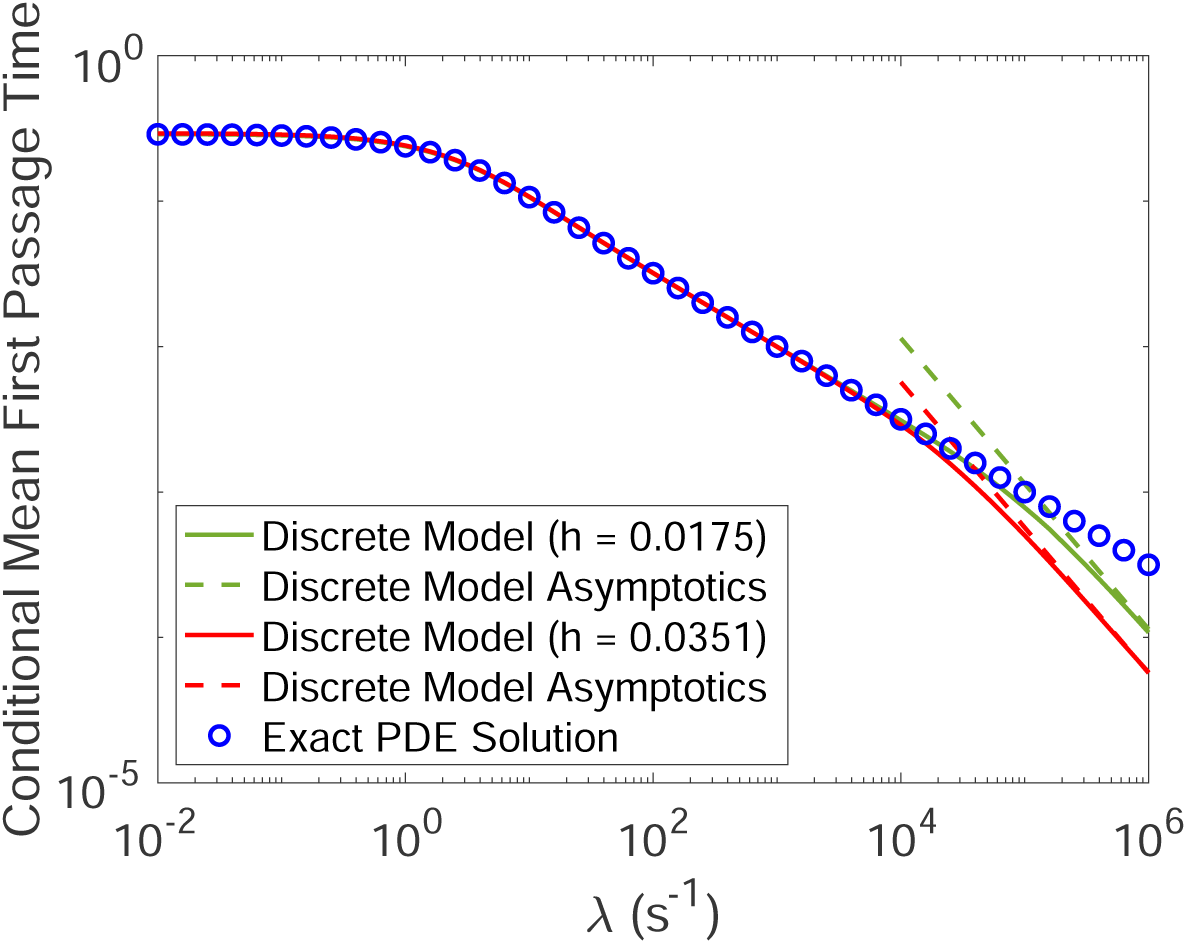
Conditional mean first passage time to reach the nucleus, ⟨*T*_*λ*_⟩ = 𝔼[*T*_*λ*_|*T*_*λ*_ < ∞], when the nucleus is a sphere of radius 3*µ*m, the cell membrane is a sphere of radius 5*µ*m, and the cytosolic space between them is open (no organelle barriers). The figure shows the exact solution ⟨*T*_*λ*_⟩ from the corresponding diffusion equation PDE (blue spheres), obtained by taking the logarithmic derivative of *F*_*λ*,0_(*R*), see Section SI3. The solid red line gives the numerical solution ⟨*T*_*λ,h*_ ⟩ to the corresponding semi-discrete model using a Cartesian grid approximation to the cytosol with mesh spacing *h* = 0.0351 (comparable to the resolution of our B cell reconstructions). The green line gives the corresponding curve when the mesh spacing is reduced to *h* =.0175. Dashed lines give the asymptotic formula for the large *λ* behavior of ⟨*T*_*λ,h*_ ⟩ (6.3). We see the continuous and discrete models agree very well until the asymptotic behavior takes over, demonstrating the different short time behavior of the underlying diffusion equation and semi-discrete diffusion equation solutions. As the mesh is refined, we also see that the semi-discrete model approximates the analytical value well to a larger value in *λ*, reflecting that the short-time breakdown of the approximation of the PDE by the semi-discrete model occurs on a shorter time-scale as the mesh is refined. See Section SI3 for details on the analytical solution and numerical simulations.

